# Control of Ribosomal RNA Synthesis by Hematopoietic Transcription Factors

**DOI:** 10.1101/2022.01.21.476118

**Authors:** Charles Antony, Subin S. George, Justin Blum, Patrick Somers, Chelsea L. Thorsheim, Dexter J. Wu-Corts, Yuxi Ai, Long Gao, Kaosheng Lv, Michel G. Tremblay, Tom Moss, Kai Tan, Jeremy E. Wilusz, Austen R. D. Ganley, Maxim Pimkin, Vikram R. Paralkar

**Affiliations:** Division of Hematology and Oncology, Department of Medicine, University of Pennsylvania Perelman School of Medicine, Philadelphia, PA 19104, USA; Institute for Biomedical Informatics, University of Pennsylvania Perelman School of Medicine, Philadelphia, PA 19104, USA; The College of Arts and Sciences, University of Pennsylvania, Philadelphia, PA 19104, USA; Cardiovascular Institute, University of Pennsylvania Perelman School of Medicine, Philadelphia, PA 19104, USA; Biochemistry and Molecular Biophysics Graduate Group, University of Pennsylvania Perelman School of Medicine, Philadelphia, PA 19104, USA; Beijing Advanced Innovation Center for Genomics (ICG) & Biomedical Pioneering Innovation Center (BIOPIC), Peking University, Beijing 100871, China; Division of Hematology, Children’s Hospital of Philadelphia, Philadelphia, PA 19104, USA; Department of Biochemistry, School of Medicine, Southern University of Science and Technology, Shenzhen, China; Laboratory of Growth and Development, St Patrick Research Group in Basic Oncology, Cancer Division of the Quebec University Hospital Research Centre (CRCHU de Québec-Université Laval), Québec, QC, G1R 3S3, Canada; Department of Molecular Biology, Medical Biochemistry and Pathology, Laval University, Québec, QC, G1V 0A6, Canada; Department of Pediatrics, Children’s Hospital of Philadelphia, Philadelphia, PA 19104, USA; Department of Genetics, University of Pennsylvania Perelman School of Medicine, Philadelphia, PA 19104, USA; Department of Cell and Developmental Biology, University of Pennsylvania Perelman School of Medicine, Philadelphia, PA 19104, USA; Verna and Marrs McLean Department of Biochemistry and Molecular Biology, Therapeutic Innovation Center, Baylor College of Medicine, Houston, TX 77030, USA; School of Biological Sciences, University of Auckland, Auckland 0623, New Zealand; Digital Life Institute, University of Auckland, Auckland 0632, New Zealand; Cancer and Blood Disorders Center, Dana-Farber Cancer Institute and Boston Children’s Hospital, Harvard Medical School, Boston, MA 02215, USA; Abramson Family Cancer Research Institute, University of Pennsylvania Perelman School of Medicine, Philadelphia, PA 19104, USA

## Abstract

Ribosomal RNAs (rRNAs) are the most abundant cellular RNAs, and their synthesis from rDNA repeats by RNA Polymerase I accounts for the bulk of all transcription. Despite substantial variation in rRNA transcription rates across cell types, little is known about cell-type-specific factors that bind rDNA and regulate rRNA transcription to meet tissue-specific needs. Using hematopoiesis as a model system, we mapped about 2200 ChIP-Seq datasets for 250 transcription factors (TFs) and chromatin proteins to human and mouse rDNA, and identified robust binding of multiple TF families to canonical TF motifs on rDNA. Using a 47S-FISH-Flow assay developed for nascent rRNA quantification, we demonstrated that targeted degradation of CEBPA (C/EBP alpha), a critical hematopoietic TF with conserved rDNA binding, caused rapid reduction in rRNA transcription due to reduced Pol I occupancy. Our work identifies numerous potential rRNA regulators, and provides a template for dissection of TF roles in rRNA transcription.

**HIGHLIGHTS:**

- Multiple cell-type-specific transcription factors (TFs) bind canonical motifs on rDNA.
- The hematopoietic TF CEBPA binds to active rDNA alleles at a conserved site.
- CEBPA promotes Polymerase I occupancy and rRNA transcription in myeloid progenitors.
- We present ‘47S-FISH-Flow,’ a sensitive assay to quantify nascent rRNA.

## INTRODUCTION

Ribosomal RNAs comprise over 80% of total cellular RNA, and their transcription from rDNA repeats is the most energy intensive transcriptional process in the cell (Miller and Beatty, 1969; Pederson, 2011). Mammalian cells contain several hundred copies of near-identical rDNA repeats, arranged in tandem arrays distributed across multiple chromosomes (Long and Dawid, 1980). In any given cell, a subset of rDNA repeats is activated by occupancy of the nucleolar factor UBTF and transcribed by RNA Polymerase I (Pol I) into 47S precursor rRNA (47S pre-rRNA), which is processed to mature 18S, 5.8S, 28S rRNAs and packaged with ribosomal proteins and 5S rRNA to form ribosome subunits (Engel et al., 2018; Sharifi and Bierhoff, 2018). The transcription of 47S pre-rRNA accounts for the bulk of cellular transcription (Moss et al., 2007), and its rate varies greatly between different cell types in complex multicellular organisms, including in many malignancies (Brombin et al., 2015; Hayashi et al., 2014; Hein et al., 2013; Jarzebowski et al., 2018; Pelletier et al., 2018; Poortinga et al., 2004; Zhang et al., 2014). This variation is believed to result from modulation of Pol I activity or epigenetic silencing of whole rDNA units (McStay and Grummt, 2008; Moss et al., 2019; Sharifi and Bierhoff, 2018), and different ribosome numbers are thought to be required in different cell types for their particular proteome requirements (Mills and Green, 2017). Despite the profound energy demands of ribosome biogenesis and its centrality to cellular function, relatively little is known about how the rate of rRNA transcription is fine-tuned to meet tissue-specific needs, or how malignant cells upregulate rRNA transcription to support their rapid proliferation.

Cell-type-specific transcriptomes are primarily orchestrated by transcription factors (TFs), which bind DNA at specific motif sequences and exert downstream effects (Lambert et al., 2018). The roles of TFs have been extensively studied at promoters and enhancers of Pol II-transcribed genes, where they execute multiple functions including displacing nucleosomes, recruiting chromatin remodelers and histone modifiers, facilitating chromatin looping, and recruiting Pol II and associated cofactors or inhibitors to modulate transcription. Over a thousand TFs, belonging to dozens of families, are annotated in the human genome, many with tissue-specific expression patterns and functions (Lambert et al., 2018). Some cell-type-specific TFs have been reported to localize to the nucleolus or occupy rDNA (Cai et al., 2015; Müller et al., 2010; Pande et al., 2009; Young et al., 2007; Zentner et al., 2011, 2014), however, the extent to which these TFs directly regulate rRNA transcription is unclear. The current gaps in defining the roles of tissue-specific TFs at rDNA are compounded by the inability of standard bioinformatic pipelines to readily yield rDNA-mapping signal from high-throughput datasets. As a consequence, in contrast to the large-scale dissection that has been performed of coding gene regulators across normal and malignant tissues, it remains largely unknown how cell-type-specific regulation of rRNA, the most abundant RNA in the cell, is achieved.

Hematopoiesis is the process through which hematopoietic stem cells undergo hierarchical differentiation and maturation into diverse blood cell lineages that are essential for health and survival (Rieger and Schroeder, 2012). Over the past decades, the TFs and chromatin factors regulating hematopoiesis have been mapped in detail, making it a paradigmatic system for the study of lineage-specific transcriptional regulation (Cedar and Bergman, 2011; Liggett and Sankaran, 2020). Over 10-fold variation in rRNA transcription is observed across different normal hematopoietic progenitors and mature cells (Hayashi et al., 2014; Jarzebowski et al., 2018). In the context of malignancy, acute leukemia blast cells have long been identified by their characteristic prominent nucleoli (Smetana, 2009), and targeting ribosome biogenesis through inhibition of Pol I is currently being investigated in clinical trials for hematological malignancies (Khot et al., 2019). The differential regulation of rRNA during both normal development and in malignancy, and the therapeutic relevance of discovering cell-type-specific regulators of Pol I, thus makes hematopoiesis an ideal model system for investigating TF roles in the regulation of rRNA transcription.

In this work, we generate an atlas of TF-rDNA binding in mammalian hematopoiesis. Through customized mapping of ∼2200 publicly available ChIP-Seq datasets for ∼250 TFs and chromatin proteins to human and mouse rDNA, we identify robust, high-confidence patterns of rDNA occupancy for numerous key hematopoietic TFs to their canonical motif sequences. To demonstrate the ability of our atlas to identify functional TF binding, we focus on CEBPA, an essential TF for myeloid lineage hematopoiesis, and demonstrate its binding to actively transcribed rDNA alleles at the specific conserved site identified through our atlas. Critically, using a degron approach that allows us to assess the immediate consequences of endogenous CEBPA degradation in a physiologically relevant mouse myeloid cell line, and using a precise nascent rRNA assay that we term ‘47S-FISH-Flow,’ we find that Pol I occupancy and rRNA transcription are rapidly impaired following CEBPA degradation. Thus, our work uncovers numerous potential cell-type-specific regulators of rRNA transcription, and provides a template for dissection of TF-rDNA regulation by establishing a direct role for the myeloid TF CEBPA in rRNA transcription.

## RESULTS

### Generating an atlas of TF-rDNA binding in human and mouse hematopoiesis

To systematically interrogate whether hematopoietic TFs bind to rDNA repeats **(Fig 1A, S1)**, we first compiled from the published literature a comprehensive list of 192 TFs with reported roles in hematopoiesis, or with family members having such roles **(Table S1)**. These included TFs and TF families reported to regulate self-renewal and lineage specification, differentiation and maturation, inflammation and immunity, tumor suppression and oncogenesis, as well as TFs with hematopoietic-specific lethality in CRISPR screens (Meyers et al., 2017). We performed a manual curation of ENCODE (Davis et al., 2018) and GEO/SRAdb (Zhu et al., 2013) portals to identify publicly deposited human and mouse ChIP-Seq datasets for these TFs, as well as for 134 additional epigenetic factors, chromatin factors, and histone modifications **(Fig S2, Table S1, see Methods)**. We downloaded raw sequencing files for these experiments, from hematopoietic as well as non-hematopoietic cell types, and mapped them to custom human or mouse genome assemblies in which a single rDNA sequence of the respective species had been added **(Fig S1)** (Gonzalez and Sylvester, 1995; Grozdanov et al., 2003). Our datasets included positive controls (ChIP-Seq for core rDNA transcription machinery UBTF, TBP, Pol I) as well as negative controls (genomic input and IgG ChIP-Seq from multiple cell types). A total of 1249 human and 909 mouse datasets for 249 and 198 factors respectively (median of 4 datasets per factor per species) passed quality control metrics, forming our rDNA mapping atlas **(Fig 1B**, S2). These included 809 human and 586 mouse datasets for 155 and 123 TFs respectively. Positive and negative controls showed tracks with expected rDNA binding patterns in both species **(Fig S1)**, validating our pipeline.

**Figure 1.**
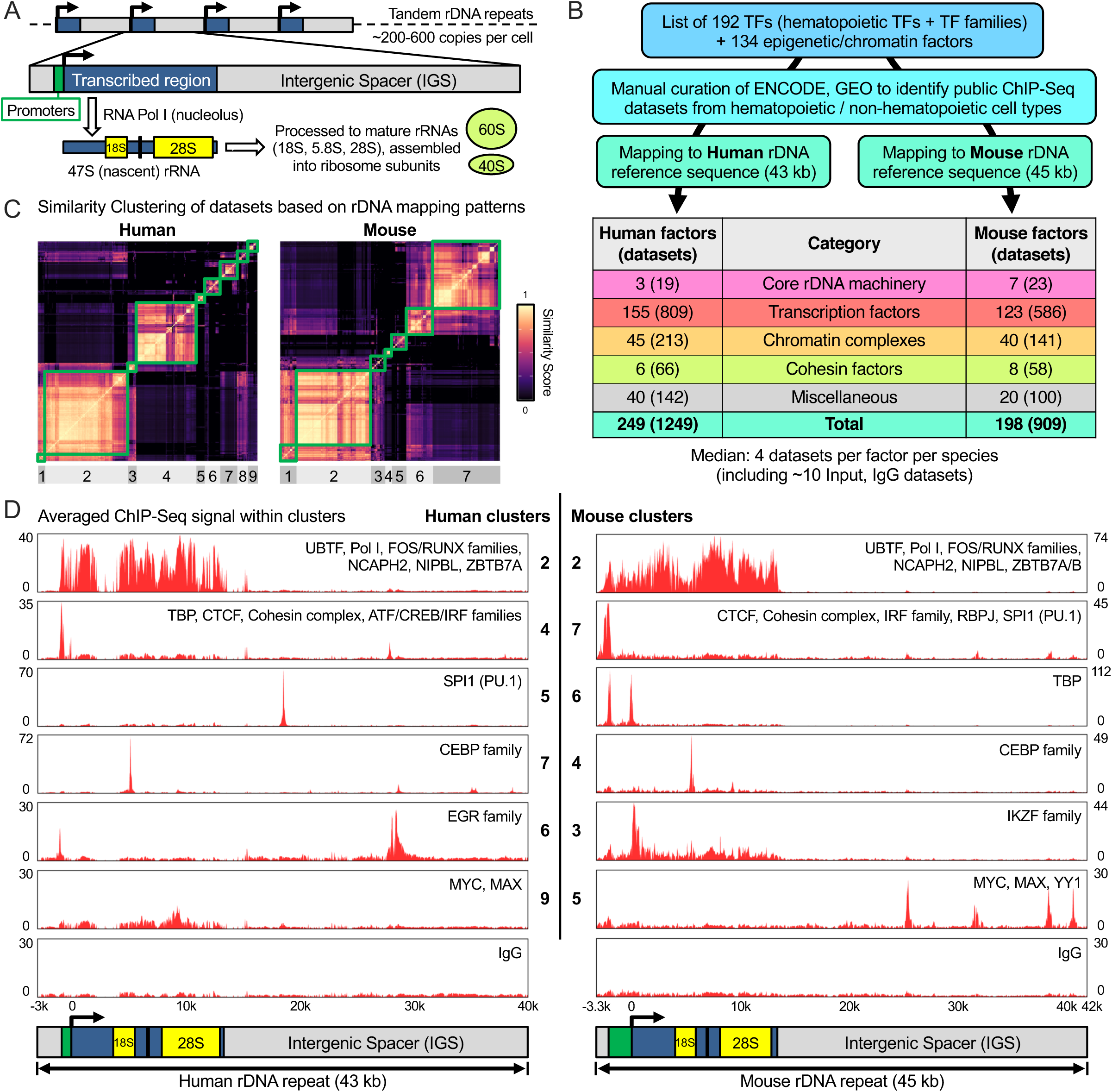
**Generating an atlas of TF-rDNA binding in human and mouse hematopoiesis** (A) rDNA organization and transcription: Every nucleated cell contains several hundred copies of rDNA repeats distributed across 5 chromosomes, a subset of which are transcribed by RNA Polymerase I into 47S (nascent) rRNA, which is processed to mature rRNAs and assembled into ribosome subunits. The length of each rDNA repeat is 43 kb in human and 45 kb in mouse. (B) Simplified pipeline for TF-rDNA atlas, involving mapping of manually curated ChIP-Seq datasets from ENCODE and GEO to human and mouse reference ribosomal DNA sequences. Number of factors represented in the atlas (with number of individual datasets in brackets) are listed for each category of factors. A detailed pipeline is provided in **Fig S2**. (C) Similarity Clustering Matrices obtained by clustering human and mouse datasets on the basis of pairwise Similarity Scores. Green squares mark individual clusters (9 in human, 7 in mouse), each representing a distinct ChIP-Seq rDNA mapping pattern. (D) Averaged rDNA ChIP-Seq mapping tracks for selected clusters, and negative control IgG, in human and mouse, showing averaged signal for all datasets within each cluster. Y-axis values depict fold-change of signal over median across rDNA length. Within each cluster, factors that met criteria for First Tier (high-confidence) rDNA binding are listed. Schematics of human and mouse rDNA are depicted at the bottom, with the green segment encompassing the rDNA Spacer promoter and 47S promoter, the blue and yellow segments marking the ∼13-kb transcribed region, and the grey segments marking the intergenic spacer (IGS). Detailed rDNA schematics are provided in **Fig S1**. See also **Fig S1, S2** and **Table S1, S2**.

**Figure 2.**
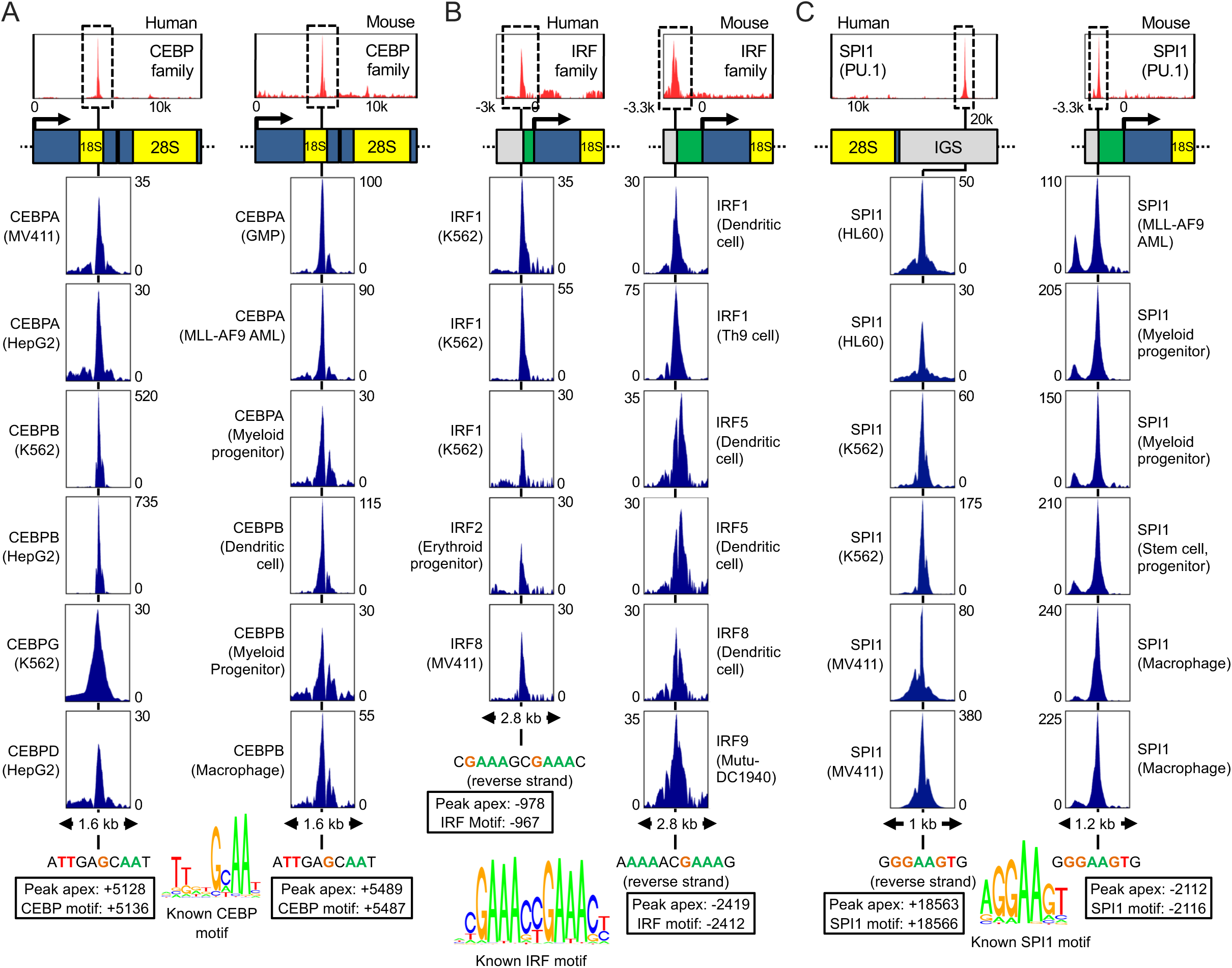
**Multiple TFs bind to canonical motif sequences on rDNA** Averaged tracks (red) of rDNA binding for (A) CEBP family members, (B) IRF family members, and (C) SPI1 in human and mouse, along with zoomed-in views of selected individual ChIP-Seq tracks (dark blue). Individual TF names are listed, with cell line or tissue type in brackets. Y-axis values depict fold-change of signal over median across rDNA length in that track. Motif sequences identified at apexes of peaks are depicted below plots, adjacent to known canonical consensus motifs for those TFs. rDNA nucleotides that match the consensus motif are highlighted with colored letters. Locations of peak apexes and motifs relative to 47S TSS in each species are boxed. See also **Fig S3, S4** and **Table S2**. Images of all First and Second Tier factor tracks are deposited to GEO.

We next aimed to interrogate our atlas for reproducible patterns of ChIP-Seq occupancy on rDNA, including identification of hubs where multiple factors might bind. Given that the atlas utilizes datasets generated by many scientific groups from diverse cell types (cell lines and primary tissues, normal and malignant) using different immunoprecipitation antibodies, ChIP-Seq protocols, and sequencing strategies, we performed global analyses in a manner agnostic to the quality or accuracy of any individual dataset. We identified all datasets showing rDNA ChIP-Seq peaks, and compared them in a pairwise fashion to each other within the same species to assign a Similarity Score between 0 and 1, reflective of similarity in locations of peaks (see Methods). Using unbiased clustering, we identified, in both species, groups of datasets that clustered together via their Similarity Scores **(Fig 1C)**. Each cluster represented a distinct pattern of ChIP-Seq signal on rDNA **(Fig 1D)**. Factors or families with at least 3 replicate datasets within a cluster (in either species) were designated as high-confidence ‘First Tier’ rDNA binding factors for that species **(Fig 1D, Table S2)**. Cognizant that polymerase-rich regions can be hotspots for non-specific ChIP-Seq signal (Teytelman et al., 2013), we applied additional stringent criteria to datasets whose signal showed broad occupancy across the entire transcribed region of rDNA, similar to Pol I occupancy (clusters 1, 2 in human and mouse); factors or families from those clusters were designated as First Tier only if 3 replicate datasets could be identified in both human and mouse. First Tier factors are listed in **Fig 1D and Table S2**, and additional lower-confidence ‘Second Tier’ factors (factors with rDNA peaks, but not meeting stringent criteria) are provided in **Table S2**. Notably, some clusters in human and mouse contained multiple factors, indicating shared or overlapping binding patterns **(Fig 1D)**.

In summary, we used public ChIP-Seq datasets relevant to the extensively-studied model systems of human and mouse hematopoiesis to generate a systematic atlas of TF-rDNA binding. This atlas defines a set of distinct TF-rDNA binding patterns, and identifies rDNA sites with binding of multiple TFs and families. We have made the atlas fully accessible by depositing to GEO (GSE193651) all mapping data, images of First and Second Tier factor tracks, as well as rDNA annotation files showing locations and sequences of peaks.

### Multiple TFs bind to canonical motif sequences on rDNA

To determine the consistency and conservation of TF binding to rDNA, we examined ChIP-Seq tracks of First Tier factors and compared binding between human and mouse. The CEBP (C/EBP, CCAAT/Enhancer Binding Protein) TF family (which has diverse roles in normal and malignant hematopoiesis (Tsukada et al., 2011)) showed consistent rDNA binding for multiple family members across multiple datasets (19 human and 13 mouse) to identical sites in human and mouse rDNA, located within the 18S rRNA region **(Fig 2A, Table S2)**, with the apex of CEBP peaks in both species precisely aligning with conserved canonical CEBP motifs. The IRF (Interferon Regulatory Factor) TF family (which plays important roles in inflammation, immunity, development, and oncogenesis (Tamura et al., 2008)) similarly showed multiple family members across multiple datasets (5 human and 15 mouse) binding to a canonical motif upstream of the rDNA promoter **(Fig 2B, Table S2)**. The IRF binding site was, in both species, located immediately upstream of previously-reported CTCF and cohesin binding sites (Herdman et al., 2017; van de Nobelen et al., 2010); we validated known CTCF and cohesin peaks through several dozen tracks in each species **(Fig S3, Table S2)**.

**Figure 3.**
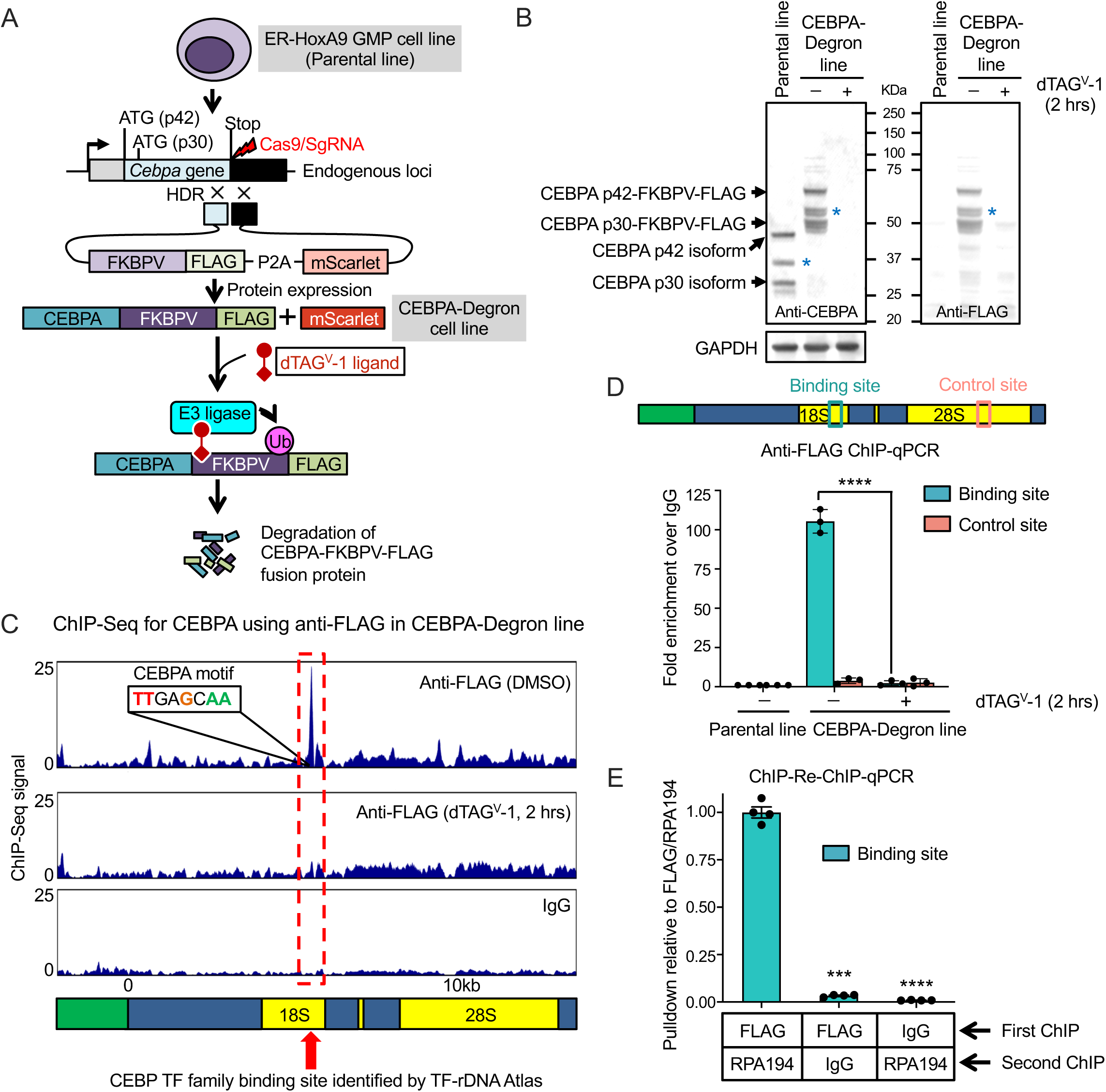
**CEBPA binds to actively transcribed rDNA repeats in a mouse myeloid progenitor line** (A) Generation of CEBPA-Degron line: CRISPR-HDR was used to integrate FKBPV degron into biallelic endogenous *Cebpa* loci in the ER-HoxA9 GMP cell line (Parental line). The resulting loci produce CEBPA-FKBPV-FLAG fusion protein, which can be ubiquitinated and degraded on treatment with dTAG^V^-1 ligand. P2A-self-cleavage site allows translation from the same mRNA of separate mScarlet protein, used to screen for clones with successful HDR. (B) Immunoblots of whole cell protein extracted from the Parental line and the CEBPA-Degron line, probed with anti-CEBPA and anti-FLAG antibodies. The Parental line shows untagged isoforms of CEBPA (p42, p30, and sumoylated p30 marked by asterisk) at their native molecular weights. The CEBPA-Degron line (without dTAG^V^-1 treatment) shows isoforms at 15 kDa larger molecular weights due to fusion of FKBPV-FLAG. The CEBPA-Degron line, on treatment with 500 nM dTAG^V^-1 for 2 hours, shows complete degradation of CEBPA. GAPDH is shown as loading control. (C) ChIP-Seq tracks in the CEBPA-Degron line using anti-FLAG antibody (pulling down CEBPA-FKBPV-FLAG fusion protein) before (DMSO) or after (dTAG^V^-1, 2 hours) degrading CEBPA, with IgG as negative control. A sharp peak is observed at the site identified in the TF-rDNA atlas (dotted red box); the CEBPA motif sequence at the apex of the peak is depicted. (D) ChIP-qPCR with anti-FLAG antibody in the Parental line, and in the CEBPA-Degron line with and without dTAG^V^-1 treatment for 2 hours. Y-axis shows fold-enrichment over IgG pulldown for each primer. n = 3 replicates. Locations of binding (cyan) and control (pale red) site primers for ChIP-qPCR are boxed on the schematic above. (E) Sequential ChIP-qPCR in the CEBPA-Degron line with anti-FLAG antibody followed by anti-RPA194 antibody (pulling down core catalytic subunit of Pol I). Pulldown combinations with polyclonal IgG are used as negative controls. Y-axis shows pulldown relative to FLAG/RPA194 samples. n = 4 replicates. All bargraphs show mean +/- standard error of mean (SEM). ***p < .001; ****p < .0001, by 1-way Anova with Sidak’s multiple comparison testing. See also **Fig S5**.

Other factors showed intriguing evolutionary divergence in binding locations between human and mouse. SPI1 (PU.1), an ETS-domain TF with crucial roles in hematopoietic stem cell self-renewal, lineage commitment, and leukemia (Antony-Debré et al., 2017; Burda et al., 2010), showed multiple datasets (8 human and 17 mouse) with abundant binding to a canonical motif in both species, but at different sites: the human SPI1 peak fell within the intergenic spacer (IGS) region, while the mouse SPI1 peak overlapped the rDNA promoter **(Fig 2C, Table S2, see also Fig S3)**. Surprisingly, MYC, long-regarded as a direct master regulator of rRNA transcription throughout the animal kingdom (Arabi et al., 2005; Grandori et al., 2005; Grewal et al., 2005; van Riggelen et al., 2010), showed rDNA binding (along with its dimerization partner MAX (Grandori et al., 2000)) at different locations in human and mouse rDNA **(Fig 1D, S4)**. Additional First Tier TFs showed binding to canonical motifs in only one species - ATF, CREB, EGR families in human, and RBPJ, YY1 in mouse **(Fig 1D, S4, see also S3)**. It is unclear whether this indicates true evolutionary divergence, or a paucity of high-quality, context-appropriate datasets for these factors in the other species.

**Figure 4.**
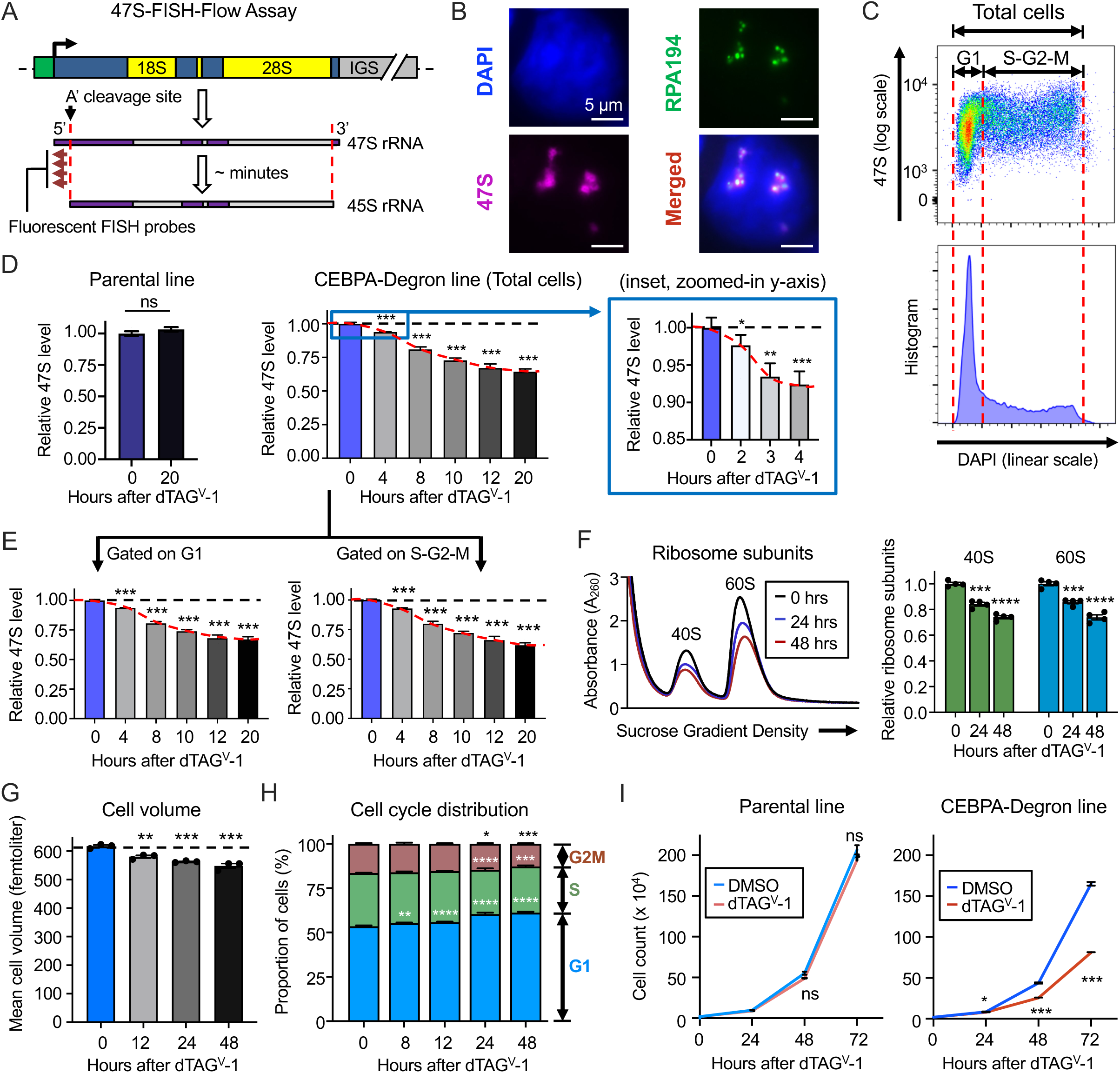
**CEBPA degradation rapidly reduces rRNA transcription** (A) Design of 47S-FISH-Flow assay: Nascent 47S pre-rRNA undergoes rapid cleavage (red dotted lines) at 5’ and 3’ ends to produce 45S rRNA. Fifteen fluorescent probes were designed to be uniquely complementary to the segment upstream of the 5’ cleavage site (A’ cleavage site) in mouse 47S pre-rRNA. (B) Widefield microscopy imaging showing nucleolar hybridization of 47S-FISH-Flow probes, along with immunofluorescence with anti-RPA194 antibody to mark actively transcribed rDNA alleles, and DAPI for overall DNA. Scale bar (white line) at bottom right indicates 5 µm. (C) Representative flow cytometry (2D scatter) plot of 47S-FISH-Flow in ER-HoxA9 cells depicting 47S FISH signal (logarithmic Y-axis) and DAPI (linear X-axis), enabling quantification of 47S pre-rRNA in different stages of the cell cycle. G1 and S-G2-M populations in the upper image are matched (using red dotted lines) to a typical DAPI histogram cell cycle profile in the lower image. (D) Left panel: Effect of dTAG^V^-1 treatment (0, 20 hrs) on median 47S level in the Parental cell line (no degron), quantified by 47S-FISH-Flow. n = 4 replicates. Middle panel: Effect of dTAG^V^-1 treatment (CEBPA degradation) on median 47S level in the CEBPA-Degron line, quantified by 47S-FISH-Flow at stipulated timepoints (0 to 20 hours) after treatment with 500 nM dTAG^V^-1. n = 4 replicates. Right panel (blue inset box): Zoomed-in Y-axis showing early time points (0 to 4 hours) following CEBPA degradation. n = 8 replicates. (E) Effect of CEBPA degradation on median 47S level in G1 and S-G2-M sub-populations in the CEBPA-Degron line, quantified by 47S-FISH-Flow. n = 4 replicates. (F) Effect of CEBPA degradation on ribosome subunit abundance in extracts prepared from identical cell numbers. Left panel: Representative A_260_ absorbance tracings. Right panel: Quantification of relative area under the curve for 40S and 60S subunits. n = 4 replicates. (G) Effect of CEBPA degradation on mean cell volume in femtoliters, measured by Coulter counter. n = 3 replicates. (H) Effect of CEBPA degradation on cell cycle distribution, gated using DAPI stain. n = 4 replicates. (I) Effect of dTAG^V^-1 treatment on growth of the Parental line (Left), and the CEBPA-Degron line (Right). n = 3 replicates. All bargraphs show mean +/- standard error of mean (SEM). *p < .05; **p < .01; ***p < .001; ****p < .0001; ns = not significant, by 2-way Anova with Sidak’s multiple comparison testing. See also **Fig S6, S7**.

Finally, several factors showed occupancy across a broad stretch of rDNA in both human and mouse, extending from the promoter throughout the transcribed region - FOS and RUNX families, ZBTB7A/B, the condensin subunit NCAPH2, and the cohesin loading factor NIPBL **(Fig 1D, S4)**. These factors matched the pattern of rDNA transcriptional machinery UBTF and Pol I **(Fig S1)**, and TF motifs were not assessed due to the absence of discrete peaks. These results are consistent with observations of RUNX family TFs interacting with UBTF and indirectly occupying rDNA chromatin (Pande et al., 2009; Young et al., 2007).

In summary, our TF-rDNA atlas allowed us to identify evolutionarily conserved as well as divergent patterns of rDNA occupancy for multiple critical hematopoietic TFs. The binding of many of these factors to discrete peaks at canonical motif sequences suggests that we have revealed direct TF-rDNA interactions of functional significance.

### CEBPA binds to actively transcribed rDNA repeats in a mouse myeloid progenitor line

We next sought to determine whether TF binding identified by our atlas is indicative of a role in rRNA regulation. We focused on the CEBP family, which belongs to the basic leucine zipper (bZIP) superfamily of TFs (Jindrich and Degnan, 2016). The CEBP family has six members (CEBPA, CEBPB, CEBPG, CEBPD, CEBPE, DDIT3) that can either homodimerize with themselves, or heterodimerize with other CEBP family members or other bZIP TFs (Rodríguez-Martínez et al., 2017; Tsukada et al., 2011). Though our atlas included datasets from a variety of bZIP TFs, no other factors showed rDNA binding patterns similar to the CEBP family **(Fig 2A, Table S2)**, indicating that the conserved motif we have identified (5136 and 5487 nt downstream of human and mouse rRNA TSS respectively) is likely bound exclusively by CEBP factors. Each CEBP TF has a characteristic pattern of expression across tissue types and within the hematopoietic tree **(Fig S5A, S5B)**, consistent with their lineage-specific and cell-type-specific roles. CEBPA (C/EBP alpha, C/EBPα), the founding member of the CEBP family, is expressed at high levels in the bone marrow granulocyte-monocyte progenitor (GMP) population **(Fig S5B)**, and its hematopoietic knockout in mice leads to loss of GMPs and downstream myeloid cells, causing lethality from infection (Pundhir et al., 2018; Zhang et al., 1997, 2004). *CEBPA* mutations are observed in 10% of human Acute Myeloid Leukemia (AML) patients (Fasan et al., 2014; Pabst et al., 2001; Papaemmanuil et al., 2016), and multiple CRISPR screens have identified CEBPA as a selective dependency in AML (Cao et al., 2021; Tsherniak et al., 2017; Tzelepis et al., 2016). Given that CEBPA has important roles in both normal and malignant myeloid biology, and that it shows conserved binding to rDNA, we sought to investigate its role in the regulation of rRNA transcription.

We mimicked physiological GMPs by using the mouse ER-HoxA9 cell line, a clonal GMP line generated by expressing Estrogen-Receptor-fused-HoxA9 in mouse bone marrow cells (Sykes et al., 2016). Culturing with beta-estradiol maintains a HoxA9-driven transcription network that keeps the line arrested in an immortalized state with flow cytometry and transcriptional profiles matching normal GMPs, but with the ability to undergo normal myeloid differentiation upon withdrawal of beta-estradiol (Blanco et al., 2021; Sykes et al., 2016). To assess the immediate effects of CEBPA loss in these GMP cells, we utilized the FKBP12^F36V^ (FKBPV) degron system, which involves fusing a target protein with the FKBPV degron domain, and treating with the small molecule dTag^V^-1 to recruit E3 ubiquitin ligase (VHL) to ubiquitinate and degrade the target (Nabet et al., 2020). The *Cebpa* gene has a single exon, with two ATG start sites that translate into two N-terminus protein isoforms (p42 and p30) with identical C-termini **(Fig 3A)**. We used CRISPR/HDR (Homology Directed Repair) in ER-HoxA9 cells to integrate an ‘FKBPV-FLAG-P2A-mScarlet’ cassette immediately upstream of the stop codon of endogenous *Cebpa* alleles, thereby fusing the FKBPV degron domain to the C-terminus of CEBPA. P2A-self-cleaved mScarlet fluorescent protein allowed selection of clones with successful HDR. We picked a bi-allelically tagged clone (hereafter referred to as ‘CEBPA-Degron line’) showing successful tagging of p42 and p30 (as well as sumoylated p30) CEBPA isoforms with FKBPV-FLAG **(Fig 3B)**. Upon treatment of the CEBPA-Degron line with dTag^V^-1, we observed complete degradation of all CEBPA isoforms within 2 hours **(Fig 3B)**. ChIP-Seq with anti-FLAG antibody confirmed binding of the CEBPA-FKBPV-FLAG fusion protein to the precise site identified by our atlas, thereby validating both that CEBPA binds its motif on rDNA, and that FKBPV fusion does not impair this binding **(Fig 3C)**. Treatment with dTAG^V^-1 for 2 hours eliminated the CEBPA peak **(Fig 3C)**. We further confirmed the specificity of binding using anti-FLAG ChIP-qPCR, and again found that pulldown of the binding site was completely lost on degradation of CEBPA with dTag^V^-1 **(Fig 3D)**. Since the CEBPA peak falls within the transcribed region of rDNA, we performed RNA immunoprecipitation (RIP-qPCR) to confirm that CEBPA does not bind ribosomal RNA and, as expected, did not find any enrichment of 47S rRNA in CEBPA pulldown **(Fig S5C)**. To assess whether CEBPA binds to rDNA copies that are being actively transcribed, we performed ChIP-Re-ChIP (Sequential ChIP), first with pulldown of crosslinked chromatin with anti-FLAG antibody, followed by elution using 3X FLAG peptide and pulldown of the eluate with anti-RPA194 antibody (RPA194 is the core catalytic subunit of Pol I). We found that when the above-mentioned antibodies were used sequentially, DNA fragments from the CEBPA binding site were pulled down at over 30-fold higher abundance than when either antibody was replaced by polyclonal IgG **(Fig 3E)**. This demonstrates that CEBPA occupies rDNA alleles that are being simultaneously transcribed by Polymerase I. In summary, CEBPA binds to actively transcribed rDNA alleles in a mouse GMP line at the site identified through our TF-rDNA atlas.

### CEBPA degradation rapidly reduces rRNA transcription

Based on our finding of conserved CEBPA binding to rDNA, we hypothesized that CEBPA regulates the transcription of rRNA. 47S pre-rRNA, the nascent rRNA transcript, is cleaved within minutes, and the abundance of 47S is therefore a good surrogate for the rate of rRNA transcription (Popov et al., 2013). To quantify 47S pre-rRNA, we employed FISH-Flow, which involves hybridizing fluorescently-labeled oligos to RNA and quantifying fluorescence using flow cytometry. This approach has previously been used to quantify intermediates of rRNA processing in hematopoietic cells (Jarzebowski et al., 2018); here we developed a modified method with probes exclusively hybridizing 47S pre-rRNA, which we term ‘47S-FISH-Flow’. We utilized the fact that 47S undergoes cleavage at defined sites at its 5’ and 3’ ends to form 45S pre-rRNA, a shorter processing intermediate on the way to mature rRNAs. The 5’ cleavage site is called the A’ site, and the segment upstream of it (650 nucleotides in mice) is unique to nascent 47S pre-rRNA **(Fig 4A, S6A)**. We designed a pool of fifteen fluorescently-labeled FISH (fluorescence in situ hybridization) probes complementary to this segment **(Fig S6A)**, performed hybridization in fixed and permeabilized cells, and confirmed with microscopy that the probes, as expected, produced bright nucleolar signal that co-localized around Pol I (RPA194) foci **(Fig 4B, S6B)**. We then used flow cytometry to quantify 47S-FISH and DAPI (DNA content) signal on a per-cell basis in the ER-HoxA9 line, giving us the ability to gate and analyze subpopulations based on cell cycle stage as needed **(Fig 4C)**. We tested the dynamic range of the assay using serum starvation, which is known to arrest rRNA transcription (Grummt et al., 1976), and could quantify rapid reduction in 47S-FISH-Flow signal that matched results seen by Northern blot **(Fig S6C, S6D)**. We also tested the assay using FKBPV-targeted degradation of RPA194 (Pol I), and found that reduction in 47S-FISH-Flow signal mirrored the reduction in rDNA-bound RPA194 (**Fig S6E-G**). Thus, 47S-FISH-Flow is able to precisely quantify nascent 47S pre-rRNA abundance and dynamics on a per-cell basis.

Since dTag^V^-1 treatment of the CEBPA-Degron line completely eliminated total and rDNA-bound CEBPA within 2 hours **(Fig 3B, 3D)**, we anticipated that any direct effect on rRNA transcription ought to be rapidly appreciable in subsequent hours. After confirming that dTag^V^-1 ligand itself had no toxic effect on 47S-pre-rRNA abundance in the Parental (no degron) cell line **(Fig 4D, left panel)**, we treated the CEBPA-Degron line with dTag^V^-1, and observed reduction in 47S levels within 4 hours, plateauing at ∼35% reduction by 12 hours **(Fig 4D, middle panel)**. Assessment of earlier time points showed that reduction of 47S abundance could be appreciated as early as 2 hours, indicating that reduction of rRNA synthesis begins almost immediately after CEBPA depletion **(Fig 4D, right panel inset)**. Because cell cycle distribution can be a confounder in rRNA quantification, and cannot be accounted for in bulk assays such as Northern blot, we exploited the ability of 47S-FISH-Flow to measure 47S in subpopulations of cells in different cell cycle stages. Separate gating and quantification from cells in G1 and in S-G2-M subsets showed that CEBPA loss equally hampered nascent rRNA levels irrespective of cell cycle distribution **(Fig 4E)**. Nucleolar size was also reduced following CEBPA degradation **(Fig S7A)**. Since doubling of ribosome mass is required for each cell division, we reasoned that a 35% reduction in rRNA transcription should be sufficient to cause reduction in mature ribosome subunit abundance. To test this, we prepared whole cell extracts from identical cell numbers at different time points after CEBPA degradation, dissociated all ribosomes into free 40S and 60S subunits using EDTA, and performed sucrose gradient centrifugation profiling to quantify subunit abundance. Consistent with the long half-lives of mature ribosomes (Hirsch and Hiatt, 1966), we observed progressive reduction (over 20%) in 40S and 60S subunits at 48 hours following CEBPA degradation **(Fig 4F)**. In parallel, CEBPA degradation also led to a slight reduction (9%) in mean cell volume **(Fig 4G)**. Since a cell needs to achieve a certain threshold of total protein content during G1 to trigger transition through the G1-S checkpoint (Schmoller et al., 2015; Zatulovskiy et al., 2020), and since impairments of ribosome biogenesis typically cause accumulation of cells in G1 (Polymenis and Aramayo, 2015), we assessed cell cycle distribution, and observed that CEBPA depletion was followed by increased proportions of cells in G1 **(Fig 4H)**. Collectively, while dTag^V^-1 ligand has no effect on the growth of the Parental line, degradation of CEBPA in the CEBPA-Degron line led to reduced growth **(Fig 4I)**. These phenotypes were not accompanied by markers of nucleolar stress like stabilization of p53 or induction of p21 (**Fig S7B, S7C**). In summary, CEBPA degradation in mouse GMPs led to reduced 47S pre-rRNA transcription within hours, followed by reduced abundance of mature ribosome subunits, accumulation of cells in G1, and reduced growth.

### CEBPA degradation reduces occupancy of Pol I and RRN3 on rDNA

Finally, we sought to identify the specific steps of rRNA transcription affected by the loss of CEBPA. Mammalian rRNA transcription involves the following sequential events **(Fig 5A)** (Engel et al., 2018; Sharifi and Bierhoff, 2018): (1) A subset of rDNA repeats in the cell is activated by occupancy of promoters and transcribed region by UBTF (detailed schematic of rDNA repeat, including Spacer and 47S promoters, is provided in **Fig S1**), (2) The SL-1 complex, comprising TBP and four TAF proteins (TAF1A/B/C/D), binds the Spacer Promoter and 47S Promoter, (3) Pol I (comprised of 13 subunits) complexed with the initiation factor RRN3 occupies the Spacer Promoter and 47S Promoter, (4) RRN3 is released from Pol I, and Pol I travels along rDNA, transcribing 47S pre-rRNA. We first quantified the whole cell abundance of these players (RPA194 as representative of Pol I, TAF1B as representative of SL-1, as well as RRN3 and UBTF), and found that none of them were changed after CEBPA degradation **(Fig 5B)**. We then used timecourse ChIP-Seq (with *Drosophila* chromatin spike-in for normalization) to quantify the occupancy of these factors on rDNA. We observed that Pol I occupancy on rDNA at the promoters and across the transcribed region was significantly reduced by 4 hours of CEBPA degradation **(Fig 5C, top left panel)**. The relative depletion of Pol I across the transcribed region at 8 hours was ∼40% **(Fig 5C, top right panel)**, concordant with the plateau of 47S pre-rRNA reduction quantified by 47S-FISH-Flow **(Fig 4D, middle panel)**. The occupancy of RRN3 was also reduced by ∼40% at the Spacer and 47S promoters **(Fig 5C, second panels)**, but, strikingly, the occupancy of SL-1 complex at promoters was unchanged **(Fig 5C, third panels)**. Similarly, UBTF did not show any reduced occupancy **(Fig 5C, bottom panels)**, and instead showed a non-significant trend towards increased occupancy. The cause of this potential increase is unclear, but may point to a compensatory mechanism occurring in response to reduced rRNA transcription.

**Figure 5.**
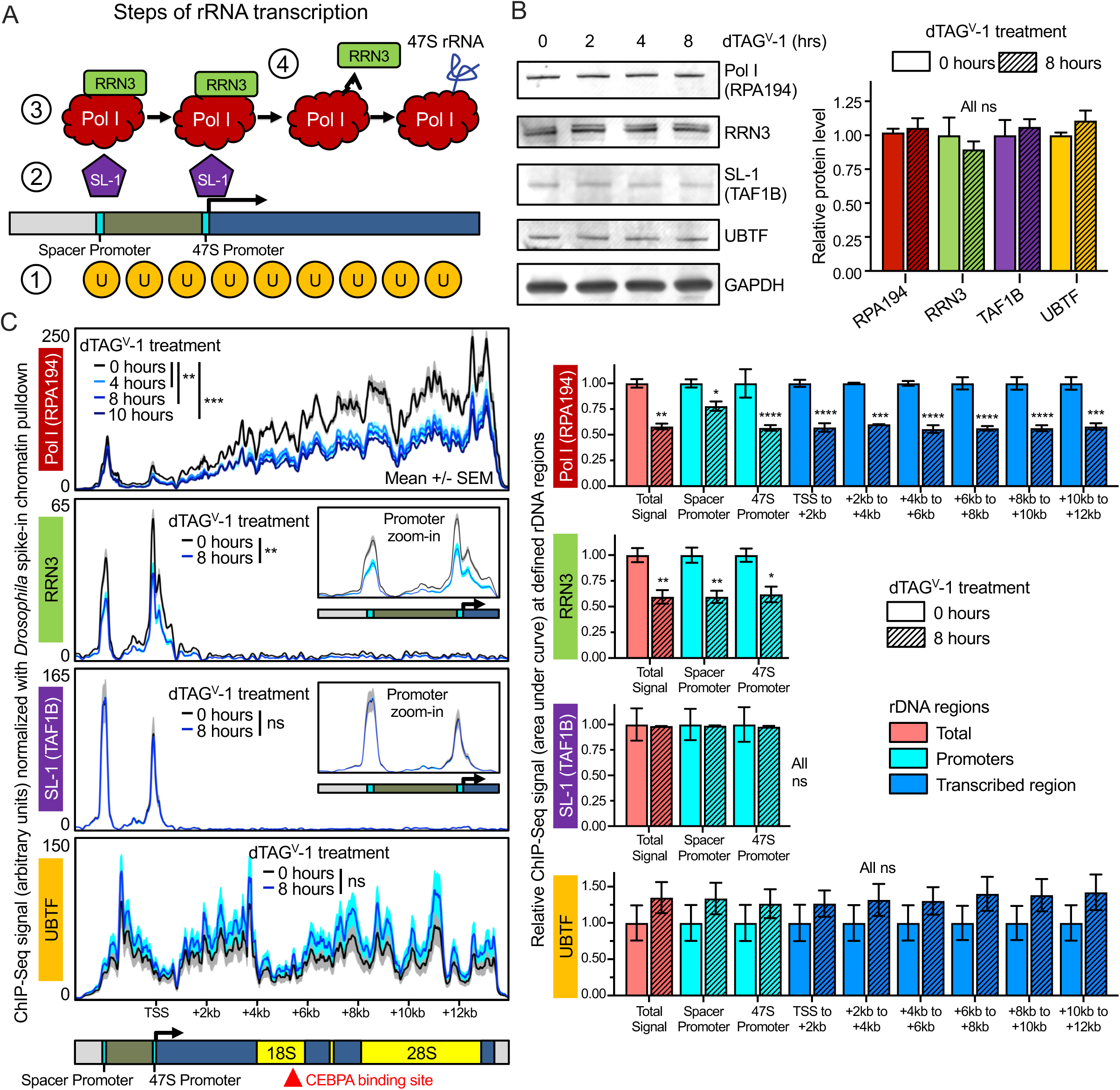
**CEBPA degradation reduces occupancy of Pol I and RRN3 on rDNA** (A) Schematic of steps involved in rRNA transcription: (1) UBTF occupancy across promoters and transcribed region, (2) Occupancy of SL-1 complex at promoters, (3) Occupancy of Pol I-RRN3 complex at promoters, (4) Detachment of RRN3, and elongation of Pol I across rDNA to transcribe 47S pre-rRNA. The schematic depicts a zoomed-in view of rDNA promoters and the immediate neighboring region, hence the CEBPA binding site falls beyond the right edge. (B) Effect of CEBPA degradation on whole cell abundance of core rDNA transcriptional machinery: Pol I (RPA194), RRN3, SL-1 complex (TAF1B), and UBTF, with GAPDH as loading control. Representative timecourse immunoblots are shown. Bargraphs show relative protein quantification at 8 hrs compared to 0 hrs, plotted as mean +/- SEM. n = 3 replicates. (C) Effect of CEBPA degradation on occupancy of core rDNA transcriptional machinery (from top to bottom) Pol I (RPA194), RRN3, SL-1 complex (TAF1B), and UBTF, quantified using ChIP-Seq signal normalized using *Drosophila* spike-in chromatin pulldown. rDNA coverage tracks are plotted in arbitrary units as mean (bold line) +/- SEM (shaded zone). For RRN3 and TAF1B, inset boxes show zoomed-in views of signal at promoters. Bargraphs show relative area under the curve (at 8 hrs compared to 0 hrs) at defined rDNA regions, plotted as mean +/- SEM. n = 2 replicates for each timepoint. *p < .05; **p < .01; ***p < .001; ****p < .0001; ns = not significant, by 2-way Anova with Sidak’s multiple comparison testing. Red arrowhead under the rDNA schematic at bottom left depicts the CEBPA binding site.

In summary, CEBPA degradation rapidly reduced occupancy of Pol I and RRN3 on rDNA without any reduction in the cellular abundance of either factor, or any reduction in occupancy of upstream machinery. This indicates that CEBPA facilitates recruitment of the Pol I/RRN3 complex to rDNA promoters.

## DISCUSSION

There is significant variation in the rate of rRNA transcription and the abundance of ribosomes across tissues, and it is believed that different cell types require different ribosome concentrations to meet their specific translational needs (Mills and Green, 2017). However, despite rRNA being the most abundant cellular RNA, the context-specific mechanisms that fine-tune its transcription in different cell types are poorly understood. Here, we showed that numerous cell-type-specific TFs show high-confidence binding to rDNA (**Fig 6**). We demonstrated this by performing a systematic survey of TF-rDNA binding in the model system of mammalian hematopoiesis, and finding canonical motif sequences at the apexes of sharp ChIP-Seq peaks for many TFs, indicating direct rDNA binding. Though our atlas was assembled with a focus on hematopoiesis, the factors and binding sites it reveals are likely to have broad significance across tissue types.

**Figure 6.**
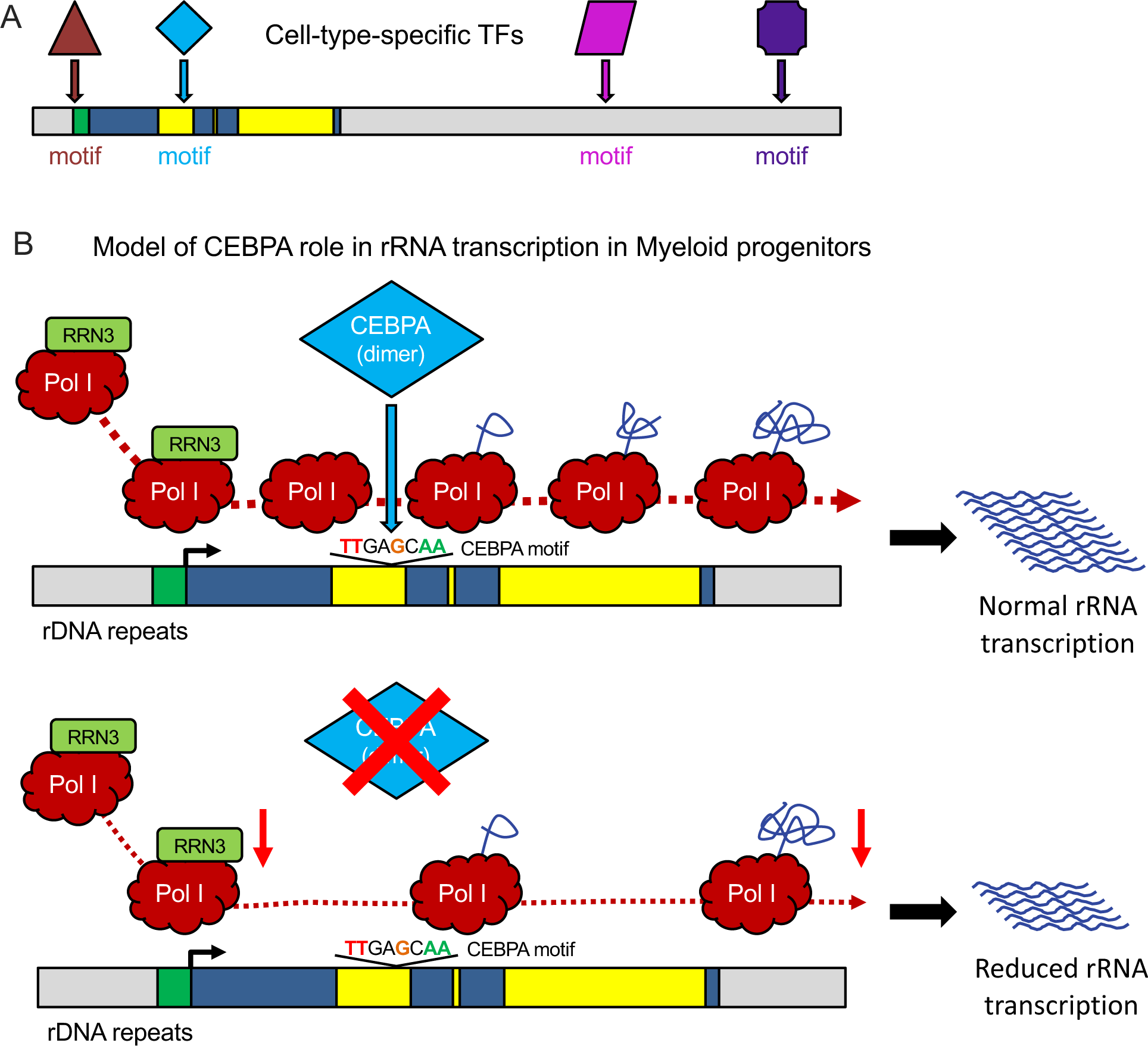
**Proposed model of cell-type-specific regulation of rRNA transcription** (A) Schematic depicting our finding that multiple cell-type-specific transcription factors bind canonical motif sequences on rDNA repeats. (B) Our model proposing that loss of CEBPA in myeloid progenitors leads to reduced recruitment of the Pol I-RRN3 complex to rDNA promoters, leading to reduced rRNA transcription.

To test the functional significance of lineage-specific TF binding to rDNA, we focused on CEBPA, a TF required for development of the myeloid lineage of hematopoiesis. An extended isoform of CEBPA (longer than the p42 isoform) has been reported to localize to the nucleolus (Müller et al., 2010), however, since no such isoform is observed in our cell system, the relevance of this prior report to our work is unclear. Our atlas identified a conserved, canonical CEBP family binding site, robustly supported by nearly three dozen independent ChIP-Seq datasets across both human and mouse. Using a cell line closely matching *in vivo* GMPs, we demonstrated CEBPA binding to actively transcribed rDNA alleles. Targeted degradation of endogenous CEBPA was followed within hours by reduced rRNA transcription, and progressively by reduced cellular ribosome numbers and impaired growth. The effect of CEBPA on rDNA appears to be to facilitate the loading of the Pol I-RRN3 complex at rDNA promoters, reflected by the reduced occupancy of these factors after CEBPA degradation **(Fig 6)**. Further investigation will be required to determine how binding of CEBPA ∼5kb downstream of the promoter mediates this effect.

The direct binding of CEBPA to mammalian rDNA, and its regulation of rRNA transcription, has implications for normal and malignant hematopoiesis. Mice with knockout of CEBPA suffer fatal loss of GMPs (Zhang et al., 2004), one of the populations with high rRNA transcription rates in the hematopoietic tree (Hayashi et al., 2014). Our work suggests that loss of CEBPA *in vivo* may lead to reduced ribosome biogenesis in GMPs, and its knockout phenotype may result partly from such an impairment. In the context of malignancy, leukemic blast cells have characteristic prominent nucleoli (Smetana, 2009), and CEBPA knockout is known to cause reduced growth in most AML lines (Tsherniak et al., 2017). We speculate that CEBPA binding to rDNA may drive high rRNA transcription rates in AML, and, if so, may serve as a cell-type-specific ribosome biogenesis vulnerability that could be exploited for treatment.

Our study relied on two critical approaches. First, we built an atlas to identify TF families with predicted functional significance in the regulation of rRNA transcription in hematopoiesis and beyond. Our atlas provides a broadly relevant resource and strong motivation to characterize the roles of the other factors whose rDNA binding we have revealed. Second, to test the functional significance of TF binding to rDNA, we used a fast-kinetics system in which depletion of a TF was coupled with timecourse measurements of polymerase occupancy and nascent rRNA. It is well-recognized that various cellular insults indirectly converge on rRNA transcription through effects on cell cycle or fitness, and studies of TF-rDNA roles that rely on overexpression or knockdown (RNAi, CRISPR) risk being confounded by an inability to distinguish direct effects from indirect ones. Approaches involving rapid depletion of endogenous TFs, such as we performed in this study, will likely be crucial. To complement fast-kinetics studies, our work also presents an assay, 47S-FISH-Flow, that provides a readout of rRNA transcription rate by quantifying the short-lived nascent 47S pre-rRNA. 47S-FISH-Flow works on a per-cell basis, can be performed on low cell numbers, has low replicate-to-replicate variability, and can quantify 47S in different stages of cell cycle. By permitting precise quantification of rRNA transcription in rare and heterogeneous populations that are otherwise challenging to assay, 47S-FISH-Flow may enable better characterization of rRNA regulation in primary tissues.

Collectively, our study opens the door to understanding how the central and essential process of rRNA transcription has been customized by evolution in complex eukaryotes to meet the ribosomal needs of different cell types, tissues, and organ systems, with implications for rRNA regulation in human disease, particularly in malignancy.

## LIMITATIONS OF THE STUDY

We assembled our TF-rDNA atlas using publicly-deposited ChIP-Seq datasets, and the atlas is therefore biased toward heavily studied factors. rDNA sequences show intra- and inter-person variations, and our use of a single reference sequence per species cannot capture the effects of variants on TF peaks. In our CEBPA studies, our degron system degrades all CEBPA protein, which, in addition to rDNA, is already known to bind tens of thousands of promoters and enhancers across the genome. Though we deem it highly likely that the rapid effects of CEBPA degradation on Pol I occupancy and rRNA transcription are direct, it is possible that delayed phenotypes on cell cycle and growth are due to a combination of impaired ribosome biogenesis as well as other transcriptome-wide changes at Pol II-transcribed genes. Uncoupling the rDNA binding of CEBPA from its genome-wide binding will be required in the future to fully distinguish these two roles.

## Supporting information

Key Resource Table

## ACKNOWLEDGEMENTS

V.R.P. is supported by National Institute of Health (NIH) grants R35-GM138035 (NIGMS) and R01-HL155144 (NHLBI), American Cancer Society (ACS) grant 129784-IRG-16-188-38-IRG, and an American Society of Hematology (ASH) Faculty Scholar Award. Y.A. is supported by the American Heart Association grant 827222. M.G.T. and T.M. are supported by Canadian Institutes of Health Research (CIHR) grant MOP12205/PJT153266 and Natural Science and Engineering Council (NSERC) of Canada grant RGPIN-2017-06128. K.T. is supported by NIH grants R01-HD89245 (NICHD) and U01-CA226187 (NCI). J.E.W. is supported by the NIH grant R35-GM11973 (NIGMS) and is a CPRIT Scholar in Cancer Research. A.R.D.G. is supported by a grant from the New Zealand Marsden Fund 14-MAU-053. M.P. is supported by a Damon Runyon-Sohn Pediatric Cancer Fellowship and the Alex’s Lemonade Stand Foundation Young Investigator Award. We thank Drs. David Sykes and David Scadden for providing the ER-HoxA9 cell line, Dr. Wei Tong for access to polysome profiling equipment, Dr. Vijay Bhoj for access to a Coulter counter, the Department of Medicine in the University of Pennsylvania Perelman School of Medicine for high-throughput sequencing resources, and the Abramson Family Cancer Research Institute for equipment access. We thank Drs. Arjun Raj, Gerd Blobel, Nancy Speck, Ivan Maillard, Peter Klein, Olivier Gadal, Konstantin Panov, and all members of the Paralkar laboratory for valuable scientific and technical discussions.

## AUTHOR CONTRIBUTIONS

V.R.P. conceived the project and supervised the study. C.A. and V.R.P wrote the manuscript with input from all authors, with M.P and A.R.D.G. providing detailed conceptual and scientific input. C.A. designed and performed all experiments relevant to CEBPA. V.R.P. curated public datasets for the TF-rDNA atlas and designed bioinformatic and analytical pipelines. L.G. and K.T. provided customized genomes for rDNA mapping. M.P. contributed ChIP-seq datasets for mapping optimization. S.S.G. performed ChIP-Seq mapping and rDNA data extraction, and J.B. wrote scripts for downstream analysis and graphical representation. D.J.W-C contributed to script writing for graphical representation. P.S. and C.A. designed and optimized the 47S-FISH-Flow assay. C.L.T. assisted in generation of the CEBPA-Degron line, and K.L. assisted in ribosome subunit profiling. M.G.T. and T.M. provided RRN3 and TAF1B antibodies and advice on their use. Y.A. performed Northern blotting under the supervision of J.E.W.

## STAR METHODS

## RESOURCE AVAILABILITY

### Lead Contact

Further information and requests for resources and reagents should be directed to and will be fulfilled by the Lead Contact, Vikram R. Paralkar

### Materials Availability

Cell lines and plasmids generated in this study are available upon request.

### Data and Code Availability

- All datasets generated in this work are deposited to GEO under the SuperSeries GSE193651: (1) SubSeries GSE191272 containing (a) Spreadsheets with rDNA mapping signal for all Human and Mouse atlas datasets, (b) Images of all Human and Mouse First and Second Tier factor tracks, and (c) Snapgene files with Human and Mouse rDNA annotations of First Tier TF peaks and motifs, and (2) SubSeries GSE193569 containing all ChIP-Seq datasets generated in the mouse CEBPA-Degron line.
- Original western blot and microscopy images have been deposited to Mendeley and are publicly available as of the date of publication.

## EXPERIMENTAL MODEL AND SUBJECT DETAILS

### Cell lines

The ER-HoxA9 cell line was a kind gift from Drs. David Sykes and David Scadden at Harvard Medical School. Chinese Hamster Ovary (CHO) cell line that stably secretes SCF was a kind gift from Dr. Andres Blanco at the University of Pennsylvania.

## METHOD DETAILS

### Generation of TF-rDNA Atlas: a) Compilation of Transcription Factor and Chromatin Protein list

A list of 192 transcription factors (TFs) was compiled from a review of the published literature using the following criteria: (1) TFs with published roles in hematopoietic stem, progenitor, differentiated, or malignant cells of any lineage, (2) All members of the following TF families/groups (AP-1, ATF, CEBP, CREB, EBF, EGR, ETS, GATA, HIF, IRF, MAML, MEF2, MYC/MAX/MAD, NFKB, NOTCH, PAX, RUNX, STAT, TCF), (3) TFs with hematopoietic-cell-type-specific dependencies (ie: selective dependencies in one or more of ALCL, AML, B-ALL, Burkitt Lymphoma, CML, DLBCL, T-ALL, or T cell lymphoma cell types) in the Broad Institute DepMap CRISPR screen (DepMap 19Q3 public release) (Meyers et al., 2017), (4) TFs mutated in hematopoietic malignancies. A list of 134 additional factors were included belonging to the following groups: (1) Known core rDNA transcription machinery, (2) Chromatin binders/readers/writers, (3) Chromatin remodelers, (3) Polymerase II factors and associated factors, (4) Cohesin factors, (5) Condensin factors, (6) DNA methylation complexes, (7) Histone modifications. The complete list of factors is provided in **Table S1** and additional details are provided in **Fig S2**.

### Generation of TF-rDNA Atlas: b) Identification of publicly available ChIP-Seq datasets

Manual searches were done through the ENCODE (Davis et al., 2018), GEO/SRAdb (Zhu et al., 2013), and EMBL (Kanz et al., 2005) portals to identify publicly available human or mouse ChIP-Seq (or ChIP-exo, CUT&RUN) datasets for the factors listed in **Table S1**. Studies from hematopoietic cell types were prioritized, but datasets from non-hematopoietic cell types were included when available, with the goal of identifying up to 20 datasets for each factor. ∼10 Input and ∼10 IgG datasets from a variety of cell types in each species were also included. A total of 1682 human and 1113 mouse datasets were identified, and raw FASTQ sequencing files were downloaded. Additional details are provided in **Fig S2**.

Each dataset was assigned a Unique ID in the following format:

Target_DatasetID_ReplicatesIfAny_CustomGenomeWithrDNA_Source_CellType. Example: A human CEBPA ChIP-Seq dataset from MV411 cells, obtained from GEO SRX2245499 (containing only 1 replicate), which we subsequently mapped to the custom hg19_rDNA genome (see below), is assigned the Unique ID: CEBPA_SRX2245499_r_hg19_rDNA_GEO_MV411. All raw FASTQ files can be obtained directly from ENCODE, GEO/SRAdb, or EMBL, depending on the Source identifier portion of the Unique ID. All datasets with Source identifier annotation ‘Pimkin’ were unpublished at time of initial mapping, but are now accessible from GEO/SRAdb under accession PRJNA751732.

### Generation of TF-rDNA Atlas: c) Data processing and rDNA mapping

Custom human and mouse genomes (which we called ‘hg19_rDNA’ and ‘mm9_rDNA’) containing a single copy of the rDNA repeat unit sequence were generated using a previously published method (Zentner et al., 2011, 2014). The human genome was the hg19 genome with a single 42,999 nt human rDNA repeat unit sequence (NCBI U13369.1 (Gonzalez and Sylvester, 1995)) inserted into chr13:15688751-15731749. The mouse genome was the mm9 genome with a single 45,306 nt mouse rDNA repeat unit sequence (NCBI BK000964.3 (Grozdanov et al., 2003)) inserted into chr12:2501-47806. With recent reports of long-read rDNA sequencing, human rDNA variants have been identified such that a single human chromosome may contain rDNA variants with single nucleotide polymorphisms or with small additional segments (Kim et al., 2018, 2021). Cognizant of this variation, we used for our analysis the reference rDNA repeat unit sequences that have been used most extensively in the literature. Detailed schematics of reference rDNA sequences are shown in **Fig S1**. Custom genomes will be provided to researchers on request.

ChIP-Seq FASTQ files were trimmed using Trimmomatic (Bolger et al., 2014) with the following parameters: *LEADING:3 TRAILING:3 SLIDINGWINDOW:4:15 MINLEN:30*, and mapped to the relevant genomes using Bowtie2 (Langmead and Salzberg, 2012) with the following parameter: *-X 2000*. SAM files from Bowtie2 were converted to BAM using Samtools View (Li et al., 2009) with the following parameters: *-F 4 -q 0*, the goal being to retain multi-mapping reads given that the genome contains numerous scattered fragments of rDNA sequences where reads of interest would map. BAM files were sorted using Samtools Sort, and indexed using Samtools Index. The IGVtools (Robinson et al., 2011) ‘count’ feature was used to extract read density across the rDNA sequence locus from the BAM files. Due to the known variability of rDNA gene copy numbers between individuals of the same species (Parks et al., 2018), as well as inconsistent availability of paired genomic input datasets for each of the downloaded ChIP-Seq files, usual normalization strategies (normalization to read number or to paired input control) could not be used. Instead, a universal normalization strategy was developed to internally normalize signal within each dataset: The read density at every nucleotide in the rDNA sequence was normalized to the median signal across the rDNA sequence for that dataset, thereby setting median coverage across each rDNA track to 1.

Additional details for data processing and mapping are provided in **Fig S2**. Generation of TF-rDNA Atlas: d) Quality assessment (QC) filtering

Datasets were retained for further analysis only if they met the following four QC criteria: (1) Trimmed read lengths longer than 30 nucleotides, (2) At least 3 million reads mapping genome-wide (rDNA repeats are present in hundreds of copies in the genome, and we empirically observed that ChIP-Seq datasets with lower total read numbers that would not have sufficed for peaks at typical alleles could nonetheless yield robust signal at the rDNA sequence), (3) At least 300K reads mapping within genome-wide peaks, as determined by MACS (Zhang et al., 2008) peak calling (this exclusion filter was not applied to negative controls [Input, IgG] or positive rDNA controls [Core rDNA transcription machinery], since they were not expected to have genome-wide peaks), (4) Non-zero read coverage across at least 75% of rDNA sequence. A total of 1249 human (for 249 factors) and 909 mouse datasets (for 198 factors) passed QC criteria. Numbers of datasets for each factor passing QC are listed in **Table S1**. Additional QC details are provided in **Fig S2**. All rDNA tracks in this manuscript were plotted using the R statistical software and the ggplot2 package (R Core Team, 2013; Wickham, 2009). Normalized rDNA mapping signal spreadsheet for all human and mouse datasets that passed QC are deposited to GEO.

### Generation of TF-rDNA Atlas: e) Similarity Score calculation and Clustering

Due to the known variation in rDNA gene copy number between individuals, the unavailability of paired genomic input dataset for each downloaded ChIP-Seq file, and higher absolute background signal in the rDNA gene sequence relative to the rest of the genome, conventional peak calling softwares (eg: MACS) could not be used to call rDNA peaks in our analysis. Instead, a custom script was written in the statistical software R (R Core Team, 2013) to calculate a Background Threshold for each dataset track (analysis restricted only to the 42,999nt/45,306nt rDNA repeat unit sequence in human/mouse respectively). First, the normalized signal across each track was sorted and log_10_-transformed. Because the raw signal (read density in each track) had been normalized by setting the median as 1, the median signal for the sorted-log_10_-transformed string for each dataset was therefore 0. An assumption was made that rDNA-binding factors of interest would not bind more than 50% of the length of the rDNA sequence, and therefore the 0 point was assumed to fall within background noise (ie: within background ChIP-seq read coverage across the entire rDNA sequence). After sorted-log_10_-transformed strings were generated, the first derivative (slope) was calculated for each string, and the first value in the string at which the slope transitioned from negative to positive was reverse-log-transformed and denoted as the Background Threshold for that track (identifying the inflection point from background noise to peak signal). Tracks for which such an inflection point could not be identified were designated as being composed entirely of background noise. For all negative controls (∼10 Input, ∼10 IgG datasets from diverse scientific groups and cell types in both species), a Background Threshold could not be identified, demonstrating that they were entirely composed of background noise and had no potential peaks. For positive controls (core rDNA transcription machinery such as UBTF, POLR1A, and TBP), the Background Threshold was able to successfully demarcate background noise from patterns of binding that matched those previously reported in the literature (Mars et al., 2018; Moss et al., 2019).

After calculation of thresholds for each of the 1249 human and 909 mouse dataset tracks, all tracks in which a Background Threshold could not be identified were excluded from further analysis as having no rDNA peaks. A custom R script was written to parse each of the remaining tracks to identify regions (peaks) with signal over Background Threshold. Adjacent peaks with <300 nt separation between them were merged. Once peaks had been identified, all datasets with peaks were compared in a pairwise fashion to each other to quantify peak overlap (performed separately within human datasets and mouse datasets) and a Similarity Score (range 0 to 1, from no overlap to exact overlap) was calculated for each pair. The following formula was used: Similarity Score = [SimilarityWeight * (Combined nucleotide length of overlapping peak regions in Dataset A and Dataset B)] / [(Nucleotide length of peak regions unique to Dataset A) + (Nucleotide length of peak regions unique to Dataset B) + SimilarityWeight * (Combined nucleotide length of overlapping peak regions in Dataset A and Dataset B)]. The SimilarityWeight correction allowed Similarity Scores to be boosted in cases of partial overlap between peaks, and was empirically fixed at 2.5 based on calibration with positive control datasets.

For each species, pairwise Similarity Scores were arranged in a 2x2 grid, and hierarchical clustering was performed on both axes using the hclust tool (ward.D2 method) in the R statistical software (R Core Team, 2013). Cluster groups were defined using the cuttree tool, and clusters with less than 9 members were excluded, as were clusters in which the maximum averaged signal across datasets in the cluster was less than 12 (indicating clusters with overall weak signal over background). The remaining datasets were re-clustered with hclust, and final groups were defined using cuttree.

### Generation of TF-rDNA Atlas: f) Designation of First Tier and Second Tier rDNA binding factors

Datasets within individual human and mouse clusters were grouped when possible based on family identity (eg: the TFs RUNX1, RUNX2, RUNX3 were grouped as ‘RUNX family’). Factors or factor families with 3 datasets within a cluster were selected as high-confidence ‘First Tier’ factors for that species. In view of concerns that polymerase-rich stretches of DNA may be particularly prone to being pulled down non-specifically in ChIP-Seq experiments (Teytelman et al., 2013), we applied further stringent criteria for clusters 1 and 2 in human and mouse, which showed broad binding across the entire transcribed region of rDNA, similar to Pol I and UBTF ChIP-Seq binding signal. Factors or families from those clusters were selected for First Tier listing only if 3 datasets could be identified in matching clusters in both human as well as mouse. In addition, tracks that showed signal solely matching the precise locations of 18S, 5.8S, and 28S rRNAs (without mapping across the rest of the 47S transcribed region) were excluded under suspicion that the original publicly deposited datasets may have been contaminated with reads from RNA-Seq libraries. First Tier factors are listed in **Table S2**. Factors with at least 2 datasets in a cluster, but not meeting stringent criteria detailed for First Tier factors, are listed in **Table S2** as lower-confidence ‘Second Tier’ factors. Track images of all First and Second Factors are deposited to GEO.

### Generation of TF-rDNA Atlas: g) Identifying TF motifs in ChIP-Seq peaks on rDNA

Peaks were identified within individual tracks as described above. For broad peaks (clusters 1 and 2 in human and mice), discrete apexes could not be identified, and TF motif searches were not performed under the assumption that broad patterns were unlikely to reflect direct motif-driven binding to DNA, and were instead likely due to secondary interactions with core rDNA machinery like UBTF or Pol I. For tracks showing discrete peaks, the peak apex was identified, and the span of the peak was defined by the points upstream and downstream at which the signal dropped to 50% of apex. The nucleotide sequence of this span was entered into the MoLoTool (Kulakovskiy et al., 2018) website (https://molotool.autosome.ru/), and a motif of the relevant factor were determined to be present if identified within 15 nt of the apex at a statistical significance of 10^-4^. For figures in this manuscript, images of known TF motifs are taken from the ISMARA (Balwierz et al., 2014) website (https://ismara.unibas.ch/mara/). Snapgene files with detailed rDNA annotations of peak and motif locations for First Tier TFs are deposited to GEO.

### Cell culture conditions

The ER-HoxA9 cell line was cultured at 37 °C in a 5 % CO_2_ atmosphere using RPMI-1640 media (Life Technologies), supplemented with 10 % fetal bovine serum (Gemini Bio), 2 % stem cell factor [SCF] conditioned media (prepared from a Chinese hamster ovary cell line that stably secretes SCF, kind gift from Dr. Andres Blanco at the University of Pennsylvania), 0.5 µM β-Estradiol (Fisher Scientific, #MP021016562), and Penicillin/Streptomycin (Thermo Fisher Scientific, #15140122). Cell counts were performed using an Accuri C6 machine (Beckman Coulter). For serum starvation experiments, centrifuged pellets were washed in Hanks’ Balanced Salt Solution (Thermo Fisher Scientific, #14025076), and were resuspended in Hanks’ Balanced Salt Solution and incubated at 37 °C for requisite periods of time.

### 47S-FISH-Flow

Fluorescent FISH probes specific to 47S pre-rRNA were designed using the Biosearch Stellaris Probe Designer tool (https://www.biosearchtech.com/support/tools/design-software/stellaris-probe-designer). 15 different FISH probes unique to 5’ ETS of 47S rRNA (650 nucleotides, see **Fig S6A** and **Key Resource Table** for details) were selected, and custom synthesized conjugated with Quasar-670 dye (Biosearch Technologies, #SMF-1065-5). The pool of FISH probe set (5 nmol) was reconstituted in 400 µl 1X TE (10 mM Tris-Cl pH 8.0, 0.5 M EDTA) to yield a stock concentration of 12.5 uM. Dissolved probes were aliquoted in single-use aliquots and stored at -20 °C. Staining was performed using modifications on a previously published protocol (Batish et al., 2011). Briefly, 5 x 10^6^ cells were harvested and washed twice with 750 µl of PBS containing 2 mM EDTA. Pelleted cells were resuspended in 750 µl of 3.7 % formaldehyde, and incubated at 37 °C for 10 min with gentle shaking. The centrifuged pellet was washed using PBS (containing 2 mM EDTA), and the resultant pellet were resuspended in 70 % ethanol to obtain a cell density of 2.5 x 10^5^ cells per 150 µl, with the ability to store at -80 °C for months. At time of staining, 2.5 x 10^5^ cells were transferred to a 96 well plate and washed using 150 µl of FISH wash buffer (10 % formamide in 2X saline-sodium citrate (SSC) buffer [300 mM Sodium Chloride, 30 mM Sodium citrate pH 7.0]). After washing, cells were resuspended in 50 µl of Hybridization buffer (10 % Dextran sulfate, 10 % formamide in 2X SSC) containing 0.5 µM fluorescent probe pool mix. Plates were sealed using parafilm and covered with aluminum foil and incubated at 37 °C overnight. After incubation, cells were pelleted and resuspended for washing in 150 µl of FISH wash buffer, and then re-pelleted and resuspended in 150 µl of FISH wash buffer for incubation at 37 °C for 30 min. Following incubation, cells were pelleted, resuspended in 150 µl of 4′,6-diamidino -2-phenylindole [DAPI] (100 ng/ml prepared in FISH wash buffer), and incubated at 37 °C for 30 min. Finally, cells were washed in 150 µl 2X SSC buffer, and resuspended in 150 µl of 2X SSC plus 150 µl of 1X PBS. 47S-FISH and DAPI intensity per cell were acquired using an LSR Fortessa flow cytometry machine (BD Biosciences) and analyzed using the FlowJo software (BD Biosciences).

For cell cycle analysis experiments, cells in 70 % ethanol were washed once in the FISH wash buffer before directly proceeding to DAPI staining.

### Generation of CEBPA-Degron line

The CEBPA-Degron line was generated by using CRISPR-HDR to integrate an FKBPV-FLAG-mScarlet cassette into the C-termini of endogenous *Cebpa* alleles in the ER-HoxA9 line as follows: A guide RNA (SgRNA_CEBPA, see **Key Resource Table**) targeting close to the stop codon of *Cebpa* was designed using the CHOPCHOP (Labun et al., 2019) website (https://chopchop.cbu.uib.no/). Tracr RNA was purchased from Thermo Scientific. SgRNA was prepared by annealing equimolar concentration of SgRNA_CEBPA and Tracr RNA under the following conditions: 95 °C x 5 min, 95 to 78 ramp [-2 °C/Sec], 78 °C x 10 min, 78 °C to 25 °C ramp [-0.1 °C/Sec], 25 °C x 5 min. The FKBPV-FLAG-P2A-mScarlet cassette flanked by 450 bp homology arms of the *Cebpa* gene was designed *in silico*, and commercially synthesized and cloned into a pUC57-derived plasmid for use as a donor template. ER-HoxA9 cells were electroporated with sgRNA/Cas9 complex and donor plasmid, using a Neon transfection kit (Thermo Fisher Scientific, #MPK1025) as follows: SgRNA (7.5 pmol) was complexed with Cas9 (1.25 µg) in buffer R (total volume 5 µl) and incubated at room temperature (RT) for 20 min to prepare sgRNA/Cas9 complex. 10^5^ cells were harvested, washed with phosphate-buffered saline [PBS] (Gibco, Life Technologies), and centrifuged at 300 x *g* for 5 min at RT. Pelleted cells were gently resuspended in 5 µl of Buffer R, and sgRNA/Cas9 complex and 300 ng of donor plasmid were added. Electroporation was performed using a 1500 mv / 20 ms / 1 pulse through the Neon transfection system (Thermo Fisher). After electroporation, cells were immediately dispensed into 1 ml of pre-warmed media without antibiotics, and kept at 37 °C for 24 hours before transferring to media with antibiotics for 4 days. Successful HDR into at least one allele resulted in the expression of in-frame mScarlet, which was used as a selectable marker for single-cell sorting using fluorescence-activated cell sorting (MoFlo Astrios, Beckman Coulter). The resulting single-cell clones were expanded for 7 days and screened for biallelic HDR by PCR (using primers CEBPA-FKBPV screen_F and CEBPA-FKBPV screen_R, see **Key Resource Table**). The final clone was confirmed through immunoblotting for tagging of CEBPA protein, and for degradation of tagged protein on treatment with dTAG^V^-1 (Tocris, #6914, reconstituted in DMSO). For all experiments involving dTAG^V^-1 treatment, cells were seeded at a density of 2 x 10^5^ cells/ml, and maintained below a maximum concentration of 10^6^ cells/ml. Based on the requirements of individual experiments, 500 nM dTAG^V^-1 or DMSO were added to media at appropriate time points prior to cell harvest. For timecourse experiments, ‘0 hr’ samples were treated with DMSO for a period of time equal to the maximum duration of dTag^V^-1 treatment in that experiment.

### Generation of RPA194-Degron line

The RPA194-Degron line was generated using a strategy similar to CEBPA-Degron line generation described above. An mScarlet-P2A-FLAG-FKBPV cassette was integrated into the N-termini of endogenous *Rpa194* alleles in the ER-HoxA9 line using CRISPR-HDR. Briefly, a guide RNA (SgRNA_RPA194, see **Key Resource Table**) targeting close to the start codon of *Rpa194* alleles was complexed with Cas9, and electroporated into the ER-HoxA9 line along with a donor plasmid template containing 450bp homology arms flanking the mScarlet-P2A-FLAG-FKBPV degron cassette. Single-cell clones were expanded and screened for biallelic integration by PCR (using primers FKBPV-RPA194 screen_F and FKBPV-RPA194 screen_R, see **Key Resource Table**). The final clone was confirmed through immunoblotting for tagging of RPA194 protein, and for degradation of tagged protein on treatment with dTAG^V^-1.

### Immunoblotting

Approximately 5 x 10^6^ cells were centrifuged at 300 x *g* for 5 min at RT, followed by washing the cell pellet with ice-cold PBS. Pelleted cells were lysed in 50 µl lysis buffer (20 mM Tris-Cl pH 7.5, 1.5 mM MgCl_2_, 140 mM KCl, 1% Triton-X-100, 0.5 mM DTT, 1 mM PMSF, 1X protease inhibitor cocktail [Sigma-Aldrich, #P8340], 2 mM Chymostatin [Cayman Chemical Company, #15114]) for 15 min on ice, and clarified twice at 17,000 x *g* for 5 and 10 min respectively. Protein concentration of the resultant lysate was measured using a BCA protein assay kit (Thermo Fisher Scientific, #23227). 80 µg of whole cell lysate was electrophoresed in a 12 % NuPAGE gel and blotted to PVDF membrane overnight at 24 V using a wet transfer blotting module (Bio-Rad). The blotted membrane was blocked in 5 % skimmed milk, followed by probing for target proteins using appropriate primary antibodies overnight at 4 °C, and probing with fluorescent conjugated secondary antibodies at 4 °C for 1 hour before imaging using a fluorescent imaging system (LI-COR odyssey). Primary antibodies and concentrations used were: anti-CEBPA 1:500 (Cell Signaling Technology, #8178S), anti-FLAG-M2 1:1000 (Sigma, #F1804), anti-GAPDH 1:1000 (Cell Signaling Technology, #2118S), anti-RPA194 1:1000 (Santa Cruz Biotechnology, #sc-48385), anti-RRN3 1:700 (proteintech, #25918-1-AP), anti-TAF1B 1:250 (generated by Tom Moss lab), anti-UBTF 1:250 (Santa Cruz Biotechnology, #sc-13125), anti-p53 1:1000 (Santa Cruz Biotechnology, #sc-126). Secondary antibodies and concentrations used were: anti-rabbit 680RD 1:1000 (LI-COR Biosciences, #925-68073), anti-mouse 800CW 1:1000 (LI-COR Biosciences, #925-32212).

### Chromatin Immunoprecipitation

25 x 10^6^ cells were harvested, washed with PBS, resuspended in 2.5 ml of 1% formaldehyde, and incubated for 15 min at RT for chromatin crosslinking. The crosslinking reaction was quenched by the addition of 125 µl of 2.5 M Glycine (final concentration 125 mM Glycine) to the suspension, followed by incubation for 5 min at RT. Cells were pelleted and washed twice with ice cold PBS. To rupture cell and nuclear membranes, pelleted cells were snap-frozen in liquid nitrogen (30 sec) followed by thawing in 37 °C (1 min) for a total of three cycles, following which the cell pellet was stored at -80 °C. On the day of experiment, the cell pellet was thawed in ice for 30 min and cells were resuspended in 400 µl of MNase reaction mix (50 mM Tris-Cl pH 8.0, 5 mM CaCl2, 0.1 mg/ml BSA, 4000 units of micrococcal nuclease) and incubated for 6 min at 37 °C for chromatin fragmentation. The MNase reaction was quenched by addition of 10 µl of 0.5 M EGTA (12.5 mM) to the suspension, and cells were pelleted and resuspended in 600 µl Sonication buffer (1X RIPA buffer containing 1 mM DTT, 0.25 % Sarkosyl [Sigma, #61747], 1X PIC, 1 mM PMSF, 2 mM Chymostatin) and split into two 300 µl aliquots in Bioruptor tubes [Diagenode, #C30010016]. Chromatin was sheared using a Bioruptor machine (Diagenode) with two cycles of 30 sec ON and 30 sec OFF. Sheared extract was clarified by centrifugation at 17000 x *g* for 10 min at 4 °C. To the clarified extract, 5 µl (50 ng) of *Drosophila* chromatin from S2 cells (Active Motif, #53083) was added as spike-in control. To ensure quantitative accuracy of spike-in, exactly 25 x 10^6^ starting cells were used for each sample, and comparative samples were harvested and processed together. Extract was precleared by incubating overnight with 5 ug of mouse polyclonal IgG (Invitrogen, #31903) in the presence of 100 ug RNase A (Fisher scientific, #FEREN0531) at 4 °C with rotation. The next morning, extract was clarified at 12000 x *g* for 10 min, and IgG was removed with Dynabeads as follows: A desired volume of Dynabeads protein G magnetic beads (Thermo Fisher, #10004D) (80 µl for each reaction) were first mixed with 3 volumes of IP buffer (50 mM Tris-Cl pH-7.5, 150 mM Nacl, 1% triton X-100, 0.5 % NP-40, 5 mM EDTA) and settled for 3 min in a magnetic stand (Fisher scientific, #FERMR02). Beads were then resuspended in 1 volume of IP buffer and ready to use. The IgG-incubated and clarified extract was mixed with 80 µl of washed Dynabeads, and incubated for 6 hours at 4 °C with rotation. To complete preclearing, all Dynabeads were removed by magnetic separation for 3 min. To the resultant supernatant (ChIP Input Lysate), 10 µg of the desired ChIP antibody [Polyclonal IgG (Invitrogen, #31903), anti-FLAG-M2 (Sigma, #F1804), anti-RPA194 (Santa Cruz Biotechnology, #sc-48385), anti-UBTF (Santa Cruz Biotechnology, #sc-13125), anti-RRN3 (generated by Tom Moss lab), anti-TAF1B (generated by Tom Moss lab)] was added, along with 2 µg of anti-H2Av (Active Motif, #61686) [which recognizes *Drosophila*-specific histone variant H2Av in spike-in chromatin], and incubated overnight in 4 °C with rotation. After incubation, the lysate was clarified at 12000 x *g* for 10 min. The supernatant was mixed with 80 µl of Dynabeads (pre-washed with IP buffer) and incubated for 6 hours at 4 °C with rotation. After incubation, Dynabeads were settled in a magnetic stand and subjected to the following washes: i) 100 mM Tris-Cl pH 7.0 [1 wash] ii) IP buffer (50 mM Tris-Cl pH 7.5, 150 mM NaCl, 5 mM EDTA, 0.5 % NP-40, 1 % TritonX-100) [6 washes] iii) 1X TE [2 washes]. Finally, beads were resuspended in 200 µl of ChIP-DNA elution buffer (100 mM Sodium bicarbonate and 1% SDS) and incubated for 5 min at RT followed by 3 min settling in a magnetic stand. The supernatant containing eluted DNA was transferred to new microfuge tubes, and 300 mM NaCl and 20 µg RNase A were added, followed by overnight shaking at 800 rpm at 65 °C in a thermo shaker (Eppendorf). The next morning, 80 µg Proteinase K (Invitrogen, #25530049) was added, followed by 2 hours of continued shaking. 1 ml binding buffer (provided in Qiagen PCR purification kit, #28106) and 20 µl of 3 M sodium acetate pH 5.5 (Invitrogen, #AM9740) were then added to the tube, and DNA was recovered using a Qiagen PCR purification kit and eluted in 50 µl of 1X TE buffer. For Next-Generation Sequencing, all of the eluted DNA was used to construct NextSeq DNA libraries using the NEBNext Ultra II DNA library prep kit for Illumina (New England Biolabs, #E7645S). Libraries were multiplexed using NEBNext Multiplex Oligos for Illumina (New England Biolabs (NEB), #E7600S), quantified using KAPA quantification kit (Fisher Scientific, #KK4824), and sequenced on an Illumina Nextseq 500 platform. For qPCR, eluted DNA was diluted 1:50 in 1X TE buffer, and qPCR was performed using 2X Luna universal qPCR master mix (New England Biolabs, #M3003X) on a ViiA7 Real-Time PCR System (Thermo Fisher Scientific).

For ChIP-Re-ChIP (Sequential ChIP) experiments, the above protocol was used with the following modifications: The first round of ChIP was performed using either anti-FLAG-M2 antibody or Polyclonal IgG. Dynabeads following the first pulldown were resuspended in Re-ChIP elution buffer (20 mM Tris-Cl pH 7.5, 150 mM NaCl, 2 mM EDTA, 1% Triton X-100, 100 µg of 3X FLAG peptide [Sigma, #F4799]), and incubated in 4 °C for 1 hour with rotation to selectively elute FLAG-fusion protein. Dynabeads were then settled in a magnetic stand, and the supernatant was used as input for the second round of ChIP (standard protocol) using either Polyclonal IgG or anti-RPA194 antibody. Eluted DNA was diluted 1:10 for qPCR.

### RIP-qPCR

RNA immunoprecipitation was performed as described elsewhere (Yao et al., 2019) except for the following modifications: Before elution of RNA, Dynabeads were incubated with 10 units of DNase I (Thermo Fisher, #18068015) for 10 min at 37 °C to eliminate DNA-mediated interactors. Beads were then washed four times in 1 ml of High salt buffer (50 mM Tris-Cl pH-7.5, 500 mM NaCl, 0.5% sodium deoxycholate, 0.1 % Igepal, 1X PIC, 1 mM PMSF, 2 mM ribonucleoside vanadyl complex) and two times in 1 ml of RIP buffer containing 500 mM NaCl. Washed beads were resuspended in 300 µl RIP-RNA elution buffer (100 mM Tris-Cl pH 6.8, 4% SDS, 10 mM EDTA), and incubated at RT for 10 min. Beads were removed, 700 ul of TRIzol (Thermo Fisher Scientific, #15596026) was added to the eluate, RNA was extracted as per manufacturer protocol, and precipitated RNA was dissolved in 20 µl nuclease free water. One-third of the isolated RNA from each sample was treated with 1 unit of DNase I for 15 min at 25 °C, following which DNase I was inactivated with addition of 1 ul of 25 mM EDTA and incubation at 65 °C for 10 min. RNA was then reverse transcribed using a Primescript RT reagent kit (Takara Bio, #RR037A) with random hexamers, followed by qPCR using Luna Universal qPCR master mix (New England Biolabs, #M3003X). Antibodies (5 µg each) used for RIP were: Rabbit IgG (Invitrogen, #02-6102), Fibrillarin (Abcam, #ab5821), Mouse IgG (Thermo Fisher Scientific, #31903), FLAG-M2 (Sigma, #F1804).

### Immunofluorescence Imaging

Microscope slides (Fisher Scientific, #12550003) were prepared by drawing a 2 cm diameter circular staining area on the upper slide face using a hydrophobic pen (Fisher Scientific, #NC9827128), and 250 µl Poly-D-lysine (Nalgene, #343910001) was dispensed into the staining area, followed by incubation overnight at 4 °C. The next morning, Poly-D lysine was removed and the circular area was washed 3X with 300 µl of nuclease free water for 5 min each, followed by 2 washes using 300 µl of PBS. After washes, 300 µl of PBS was dispensed into the staining area, and slides were kept on a slide warmer set to 37 °C. In parallel, the desired amount of cells (1.5 x 10^5^ cells per reaction) were harvested and washed once in PBS and resuspended in PBS at a density of 10^6^ cells/ml. While maintaining slides on the warmer, PBS was removed from the staining area and 100 µl of cell suspension was dispensed in its place, ensuring the volume remained within the hydrophobic circular area. Slides were then maintained at 37 °C for 1 hour to allow live cells to adhere to the poly-D-lysine coating on the slide face (we observed that allowing cells to cool below 37 °C during this step caused arrest of rRNA transcription and dissipation of nucleolar morphology). After incubation, 100 µl of 4 % paraformaldehyde in 1X PBS was added to the cell suspension meniscus in the staining area (taking care not to let the suspension overflow the hydrophobic circle), and slides were kept at RT for 15 minutes to permit fixation of adherent cells. The adhered cells were washed three times using 300 µl of PBS for 5 min each. To permeabilize cell membrane, adhered cells were treated with 50 µl of PBS and 50 ul of 0.5 % triton X-100 at RT for 5 min, and then washed three times with PBS. Adhered cells were then incubated in 50 ul of hybridization buffer (10 % dextran sulfate, 10 % formamide prepared in 2X SSC) at RT for 1 hour, followed by washing thrice with 300 ul of PBST (PBS containing 0.2 % Tween 20) for 5 min each. 150 ul of anti RPA194 antibody (1:75) prepared in 150 ul of hybridization buffer, mixed with 2 ul of 47S rRNA FISH-flow probes, was then dispensed into the hydrophobic circle, and the slide was incubated overnight at 4 °C. The next morning, the staining area was washed four times with PBST followed by incubation in secondary anti mouse AF488 (1:2000) for 1 hour in RT. The staining area was then washed four times with PBST. DAPI (250 ng/ml) prepared in PBST was added to the staining area, and incubated for 20 min in RT followed by washing four times using 300 ul of PBST for 5 min each. Cell were air dried in RT for 30 min and mounted in 50 ul of slowfade Diamond Antifade (Thermo Fisher Scientific, #S36967) using a rectangular cover glass, #1.5 thickness (Thomas Scientific, #1217N86) and sealed using nail polish.cells were imaged using a 100 X objective in a Widefield Microscope (Leica).

For nucleolar size quantification, slides were prepared as mentioned above, except that cells were blocked in 10 % serum (Cell Signaling Technology, #5425). Antibodies used were anti-RPA194 (1:75), anti-mouse AF488 (1:2000) (Thermo Fisher Scientific, #A11017), and anti-Nucleolin conjugated to AF647 (1:100) (Abcam, #ab198580).

### Northern blotting

Northern blotting was performed as described previously (Tatomer et al., 2017). 5 x 10^6^ cells were harvested and washed with PBS. Pelleted cells were resuspended in 1 ml of TRIzol (Thermo Fisher Scientific, #15596026), and RNA was isolated using manufacturer’s instructions. The RNA pellet was resuspended in 50 µl of nuclease free water. 10 µg of RNA was electrophoresed in a 1.2 % denaturing formaldehyde agarose gel with NorthernMax reagents (Thermo Fisher Scientific). The gel was washed twice in nuclease free water (15 min each) and twice in 10X SSC (15 min each) before transferring to a Hybond-N membrane (Cytiva Life Sciences, #RPN303N) through overnight capillary transfer. After incubation, the membrane was subjected to crosslinking with 120 mJ of ultraviolet radiation per unit area using a UV cross linker (Spectrolinker XL-1000). The membrane was then transferred to a hybridization tube and incubated in 10 ml of prewarmed (50 °C) ULTRAhyb oligo buffer (Thermo Fisher Scientific, #AM8663) for 45 min at 42 °C with rotation. Simultaneously, γ-^32^P labeled 47S rRNA_Northern_5’ETS probe or 28S rRNA_Northern_Loading control probe (see **Key Resource Table**) were prepared as follows: A 20 µl labeling reaction was set up with 1.5 µl of DNA oligo [10 µM], 4 µl of [γ-^32^P] ATP (PerkinElmer, #BLU502A001MC), and 10 units of T4 polynucleotide kinase (PNK) (New England Biolabs, #M0201S) in PNK buffer, and incubated at 37 °C for 60 min. The reaction was quenched by incubating at 95 °C for 5 min. Labeled probes were purified using Illustra MicroSpin G-50 columns according to manufacturer’s instructions (Cytiva Life Sciences, #27-5330-01). 15 µl of labeled probe was incubated with the membrane at 42 °C overnight. The next day, the membrane was washed twice (30 mins each) with prewarmed 42 °C Northern wash buffer (2X SSC, 0.5 % SDS), and radioactive signal was acquired using a Typhoon 9500 scanner (GE Healthcare), followed by quantification using the ImageQuant software (GE Healthcare). The level of 47S pre-rRNA per lane was normalized to 28S rRNA.

### RNA extraction and RT-qPCR

2 x 10^6^ cells were harvested and RNA was purified using Qiagen RNeasy Mini Kit (Qiagen, #74104) using manufacturer protocol. 500 ng of RNA were treated with 1 unit of DNase I at 25 °C for 15 min and heat inactivated at 65 °C for 10 min. RNA was reverse transcribed using the Primescript RT reagent kit (Takara Bio, #RR037A) with random hexamers, followed by qPCR using Luna Universal qPCR master mix (New England Biolabs, #M3003X) using recommended manufacturer protocol.

### Ribosome subunit profiling

25 x 10^6^ cells were harvested and washed with ice cold PBS. Pelleted cells were lysed in 1 ml ribosome lysis buffer (20 mM Tris-Cl pH 7.5, 140 mM KCl, 1% Triton-X-100, 0.5 mM DTT) and incubated on ice for 10 min. Lysate was clarified by centrifugation at 17000 x *g* for 10 min, and the supernatant was treated with EDTA (final concentration 60 mM) for 30 minutes on ice to dissociate all ribosomes into free ribosomal subunits. In parallel, 7-30 % sucrose gradient (20 mM Tris-Cl pH 7.5, 140 mM KCl, 5 mM EDTA, 0.5 mM DTT) was prepared using a gradient station (Biocomp instruments). 800 µl of EDTA-treated lysate was layered on top of the sucrose gradient and ultracentrifuged at 35000 rpm for 4 hours at 4 °C in a SW41Ti rotor (Beckman). After centrifugation, gradients were analyzed using a fractionator equipped with a Triax^TM^ UV flow cell (Biocomp instruments), and the A_260_ absorbance profile was recorded. Free ribosomal 40S and 60S subunit abundances were quantified by calculating the area under the curve of corresponding peaks.

### ChIP-Seq mapping and quantification in CEBPA-Degron line

FASTQ files were mapped to the mm9_rDNA genome using Bowtie2(Langmead and Salzberg, 2012), and read density across the rDNA sequence was extracted using the IGVtools (Robinson et al., 2011) ‘count’ feature, as per the pipeline described in the ‘Data processing and rDNA mapping’ section above. Since our in-house ChIP-Seq protocol incorporated *Drosophila* chromatin spike-in and H2av pulldown for the purpose of normalization, each FASTQ file was also mapped to a partial drosophila dm3 genome file (kind gift of Brian Egan, Active Motif) comprising known H2av binding regions (Egan et al., 2016). PCR duplicates were removed using a custom pipeline, and normalization of the mouse rDNA ChIP-Seq signal was performed in a three step manner, incorporating previously-published strategies (Egan et al., 2016; Mars et al., 2018): (1) rDNA signal for each dataset was normalized to the read number mapping to the partial dm3 genome for that dataset (normalizing for sample-to-sample ChIP variability), (2) For UBTF and Pol I, rDNA signal at each nucleotide was normalized to signal strength at that nucleotide in a paired input sample (normalizing for sequencing and bioinformatic biases that produce jagged region-to-region background variations in broad tracks), and (3) rDNA signal was smoothed at each nucleotide by averaging signal over a 80nt window in either direction (smoothing signal over broad regions to allow quantitative comparison). Area under curve was calculated for defined loci, and used for calculating statistical significance in timepoint comparisons of rDNA tracks using 2-way Anova with Sidak’s multiple comparison testing.

## QUANTIFICATION AND STATISTICAL ANALYSIS

### Quantification

All western blots were quantified by densitometry analysis using ImageJ software. To quantify the size of nucleoli in **Figure S7A**, microscopic images were imported to Image J software and signals were marked using optimum threshold without saturation of signal. polygon selections were made around the flourecsnce signal around nucleolin (Red channel). For each condition, 20 cells were used to quantify the are under the polygon selection and plotted as violin plot using graphpad prism software.

### Statistical analysis

All bargraphs were plotted to show mean plus or minus SEM. Statistical significance was calculated by 2-way Anova with Sidak’s multiple comparison testing using GraphPad Prism software and the significance were mentioned as follows: *p < .05; **p < .01; ***p < .001 and ****p<0.0001. ‘n’ represents the number of replicates.

---

**Figure S1 (related to Figure 1).**
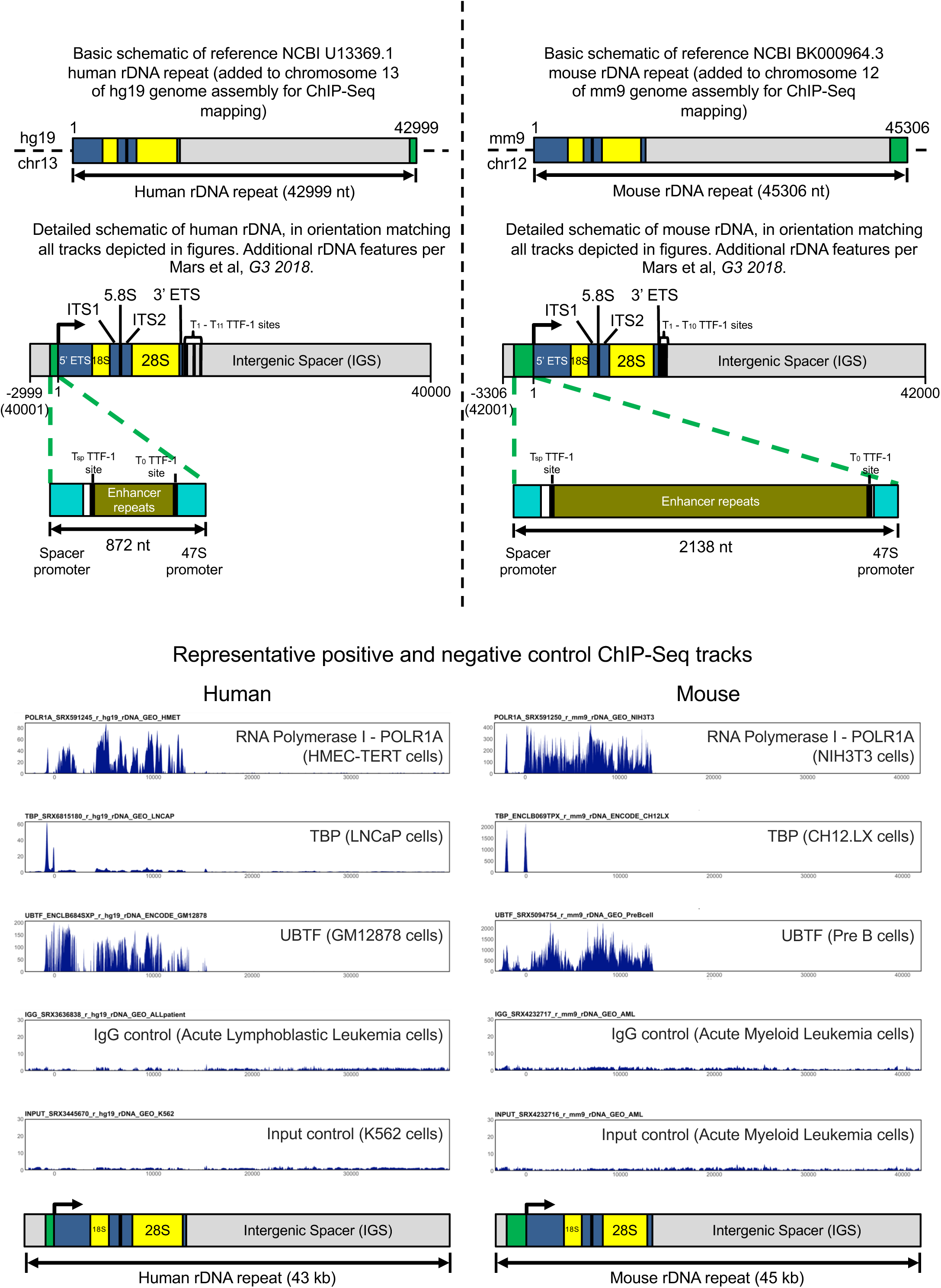
(<u>Top Left Panel</u>) Schematic of Human rDNA with annotations: The green segment encompasses the human rDNA Spacer promoter and 47S promoter, the blue and yellow segments mark the ∼13-kb transcribed region, and the grey segments mark the intergenic spacer (IGS). ITS: internal transcribed spacer; ETS: external transcribed spacer. The reference U13369.1 human rDNA repeat sequence (Gonzalez et al, *Genomics 1995*) begins at the 5’ ETS, and ends with the promoter. The unaltered U13369.1 sequence was inserted into chromosome 13 of the hg19 genome to create the custom genome used for human ChIP-Seq mapping in this paper. In all mapping tables, nucleotide 1 marks the beginning of the 5’ ETS (the TSS), and nucleotide 42999 represents the end of the 47S promoter. However, given the tandem repeat nature of rDNA, in order to better visualize contiguity of ChIP-Seq mapping between promoter and transcribed regions, the last 2999 nt stretch of U13369.1 is shown transposed upstream of TSS in all human rDNA tracks in this paper. rDNA features have been added per Mars et al, *G3 2018*. (<u>Top Right Panel</u>) Schematic of Mouse rDNA with annotations: The green segment encompasses the mouse rDNA Spacer promoter and 47S promoter, the blue and yellow segments mark the ∼13-kb transcribed region, and the grey segments mark the intergenic spacer (IGS). ITS: internal transcribed spacer; ETS: external transcribed spacer. The reference BK000964.3 mouse rDNA repeat sequence (Grozdanov et al, *Genomics 2003*) begins at the 5’ ETS, and ends with the promoter. The unaltered BK000964.3 sequence was inserted into chromosome 12 of the mm9 genome to create the custom genome used for mouse ChIP-Seq mapping in this paper. In all mapping tables, nucleotide 1 marks the beginning of the 5’ ETS (the TSS), and nucleotide 45306 represents the end of the 47S promoter. However, given the tandem repeat nature of rDNA, in order to better visualize contiguity of ChIP-Seq mapping between promoter and transcribed regions, the last 3306 nt stretch of BK000964.3 is shown transposed upstream of TSS in all mouse rDNA tracks in this paper. rDNA features have been added per Mars et al, *G3 2018*. (<u>Bottom Panels</u>) Representative example tracks of Positive controls (POLR1A, TBP, UBTF) and Negative controls (IgG, Input) mapped to Human and Mouse rDNA. UBTF and POLR1A shows expected broad binding throughout the transcribed region. TBP shows the expected two peaks at the Spacer promoter and 47S promoter. At the top left of each track is the Unique ID for the dataset, which can be used to identify specific datasets in Table S1 and in Master spreadsheets uploaded to GEO. Y-axis values are normalized to median signal across rDNA length (all tracks are shown with Y-scale of 30 or maximum signal, whichever is greater).

**Figure S2 (related to Figure 1).**
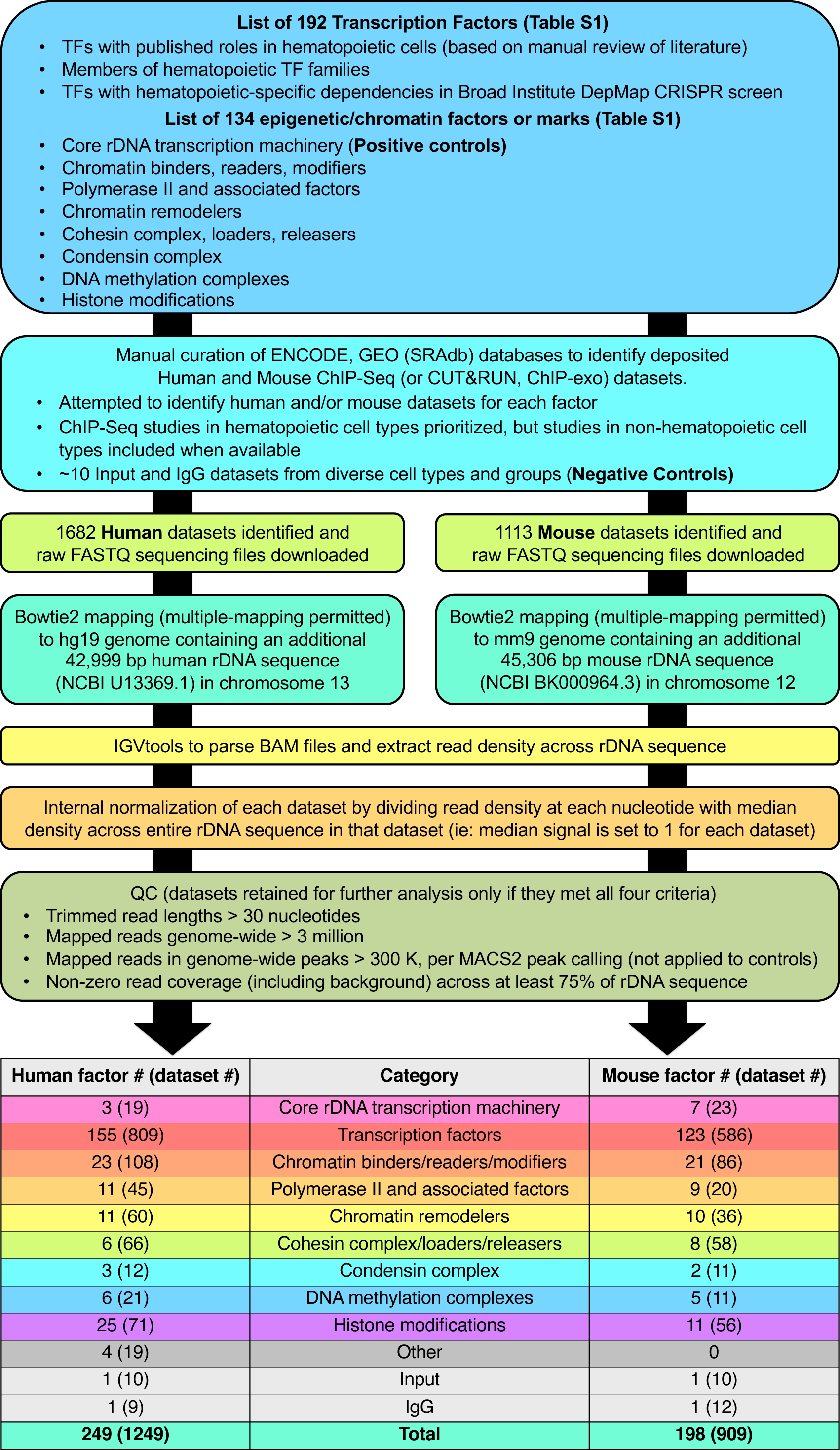
Bioinformatic pipeline for mapping of ChIP-Seq datasets from ENCODE, GEO (SRAdb) to human and mouse ribosomal DNA.

**Figure S3 (related to Figure 2).**
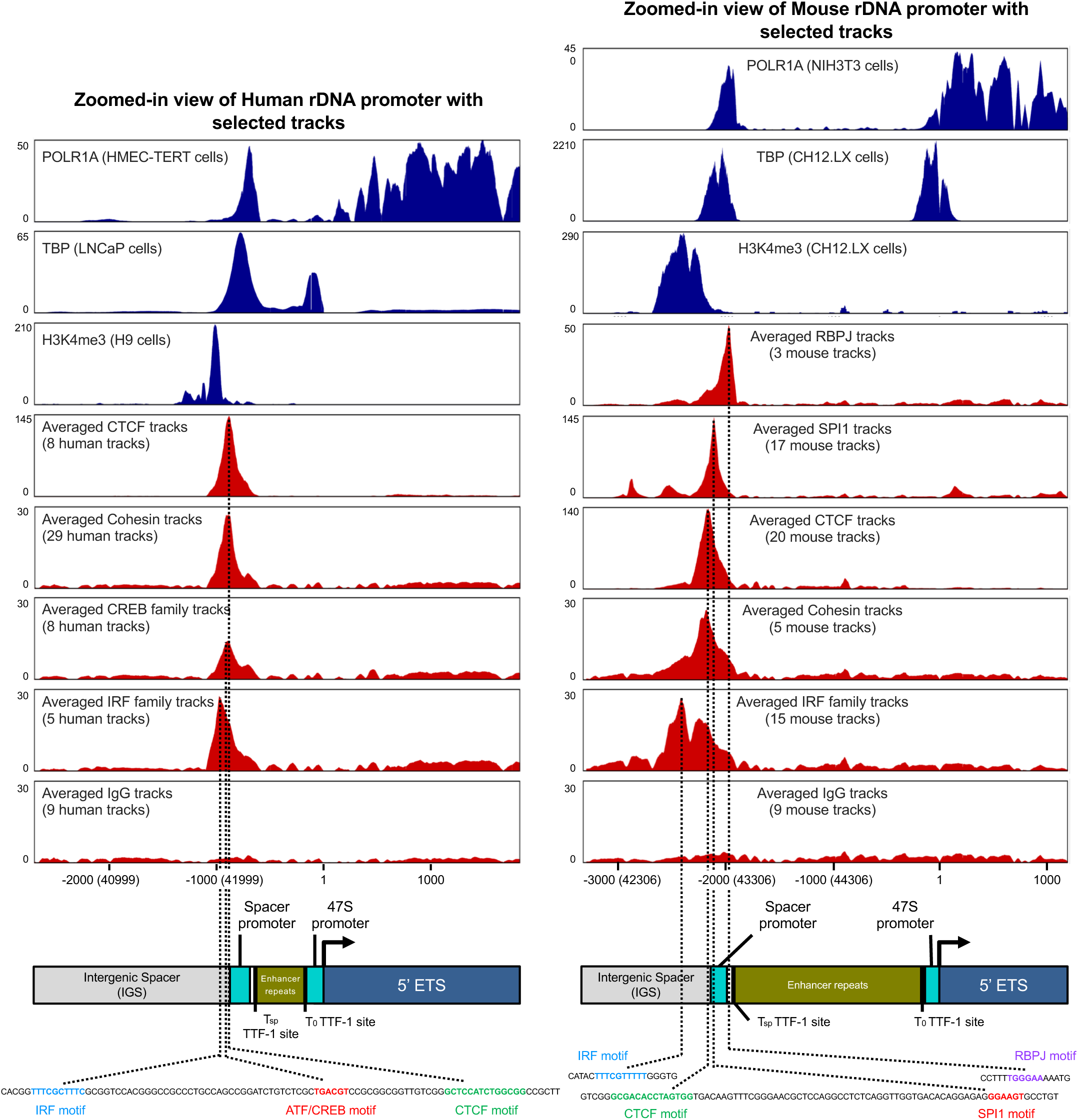
(<u>Left Panel</u>) Selected individual as well as averaged tracks (with number of factors listed) of human factors or factor families with ChIP-Seq peaks at rDNA promoter and surrounding region. The distance from the start of the annotated human Spacer Promoter to the end of the 47S Promoter is 872 nt. POLR1A binds to the Spacer Promoter, and extends from the 47S promoter into the transcribed region (starting with 5’ ETS). TBP binds to the Spacer Promoter and 47S Promoter. An H3K4me3 peak is present upstream of the Spacer Promoter. A DNA sequence stretch immediately upstream of the human Spacer Promoter is provided, containing IRF, ATF/CREB, and CTCF motifs aligning with apexes of peaks of averaged ChIP-Seq signal for the respective factors (averaged from tracks listed in Table S1). (<u>Right Panel)</u> Selected individual as well as averaged tracks (with number of factors listed) of mouse factors or factor families with ChIP-Seq peaks at rDNA promoter and surrounding region. The distance from the start of the annotated mouse Spacer Promoter to the end of the 47S Promoter is 2138 nt. POLR1A binds to the Spacer Promoter, and extends from the 47S promoter into the transcribed region (starting with 5’ ETS). TBP binds to the Spacer Promoter and 47S Promoter. An H3K4me3 peak is present upstream of the Spacer Promoter. DNA sequence stretches upstream and overlapping with the mouse Spacer Promoter are provided, containing IRF, CTCF, SPI1, and RBPJ motifs aligning with apexes of peaks of averaged ChIP-Seq signal for the respective factors (averaged from tracks listed in Table S1). Y-axis values are normalized to median signal across rDNA length (all tracks are shown with Y-scale of 30 or maximum signal, whichever is greater).

**Figure S4 (related to Figure 2).**
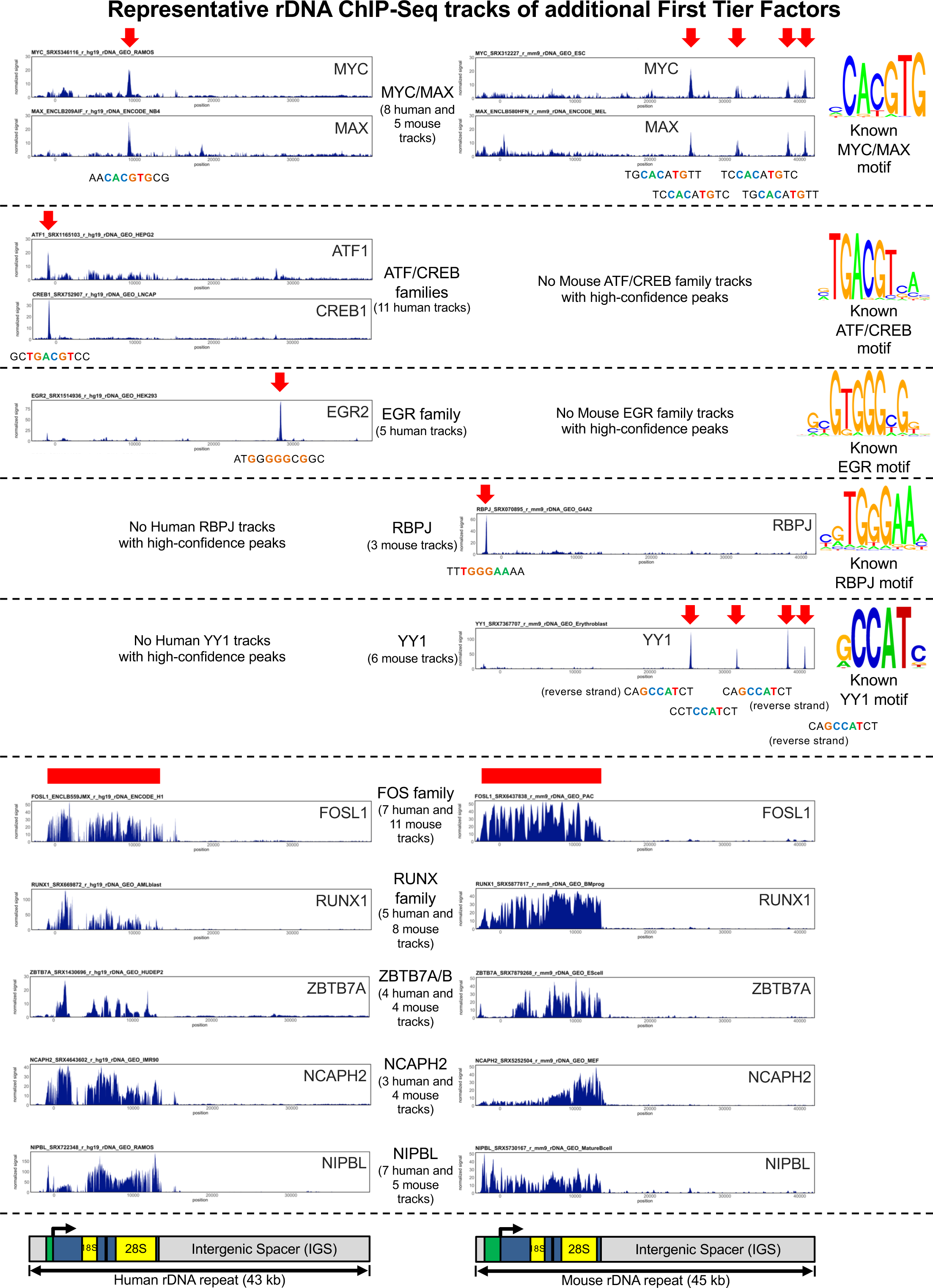
Selected Human and Mouse ChIP-Seq tracks of additional First Tier Factors binding to human or mouse rDNA (or both), with number of tracks in each species listed. Above each track is the “Unique ID” of each dataset, corresponding to Table S1 and in Master datasets deposited to GEO. Common binding sites across all tracks in a cluster are marked by red arrows. Broad ChIP-Seq signal regions are marked with a red band. A notable caveat is that the multiple consecutive MYC/MAX and YY1 peaks in the mouse IGS are at sequences known to be repeated (near-identical) within the single IGS region of the mouse reference sequence, limiting the ability to reliably distinguish between them. It is therefore unknown if these represent truly distinct binding sites. Binding motif sequences are shown for factors with discrete peaks. For broad pattern of binding without a consistent peak apex, no analysis for TF motif sequencing was performed. Y-axis values are normalized to median signal across rDNA length (all tracks are shown with Y-scale of 30 or maximum signal, whichever is greater). Motif from ISMARA (www.ismara.unibas.ch).

**Figure S5 (related to Figure 3).**
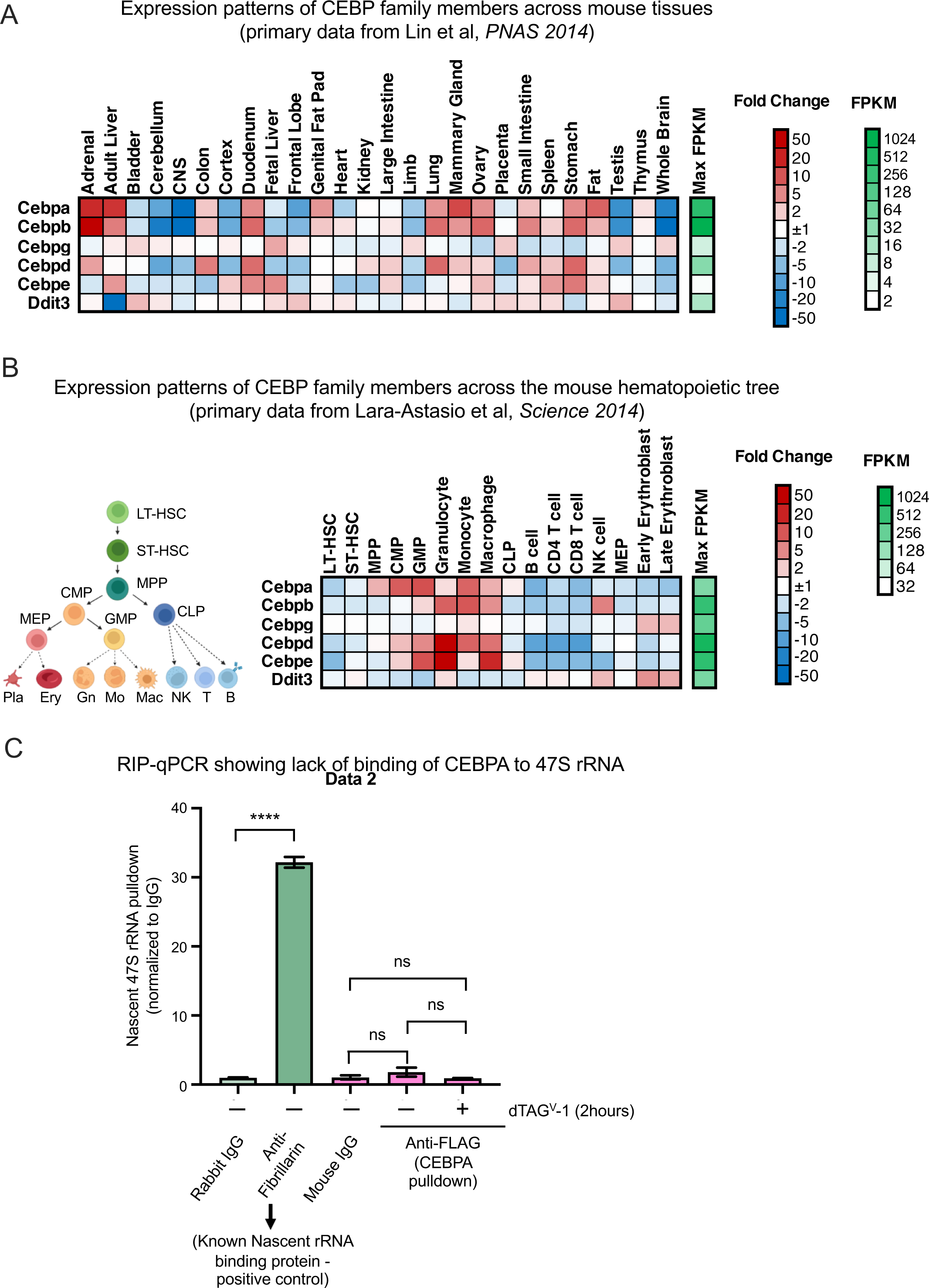
(A) Heatmap of expression patterns of the six CEBP family members across different normal mouse tissues, generated using primary data from ENCODE/CSHL Long-RNA-Seq Transcriptome project, Lin et al, *PNAS 2014*. Max FPKM (Fragments Per Kilobase of transcript per Million mapped reads) for each gene represents the RNA-Seq expression value in the tissue with the most abundant expression of that gene. (B) Heatmap of expression patterns of the six CEBP family members across the mouse hematopoietic tree, generated using primary data from Lara-Astiaso et al, *Science 2014*. FPKM = Fragments Per Kilobase of transcript per Million mapped reads. Max FPKM (Fragments Per Kilobase of transcript per Million mapped reads) for each gene represents the RNA-Seq expression value in the tissue with the most abundant expression of that gene. LT-HSC: Long-term hematopoietic stem cell, ST-HSC: Short-term hematopoietic stem cell, MPP: Multipotent progenitor, CMP: Common myeloid progenitor, GMP: Granulocyte-monocyte progenitor, CLP: Common lymphoid progenitor; MEP: Megakaryocyte-erythroid progenitor. (C) RIP-qPCR (RNA Immunoprecipitation) for nascent 47S rRNA using Anti-Fibrillarin or Anti-FLAG antibodies (with rabbit and mouse polyclonal IgG as matching negative controls) in the CEBPA-Degron line (in which endogenous CEBPA alleles have been tagged with FKBPV-FLAG). Fibrillarin, a protein known to bind nascent rRNA, shows abundant 47S pulldown and is provided as a positive control for RIP. Anti-FLAG (CEBPA) does not show enrichment over IgG in cells with intact or degraded CEBPA (-/+ dTAG^V^-1).

**Figure S6 (related to Figure 4).**
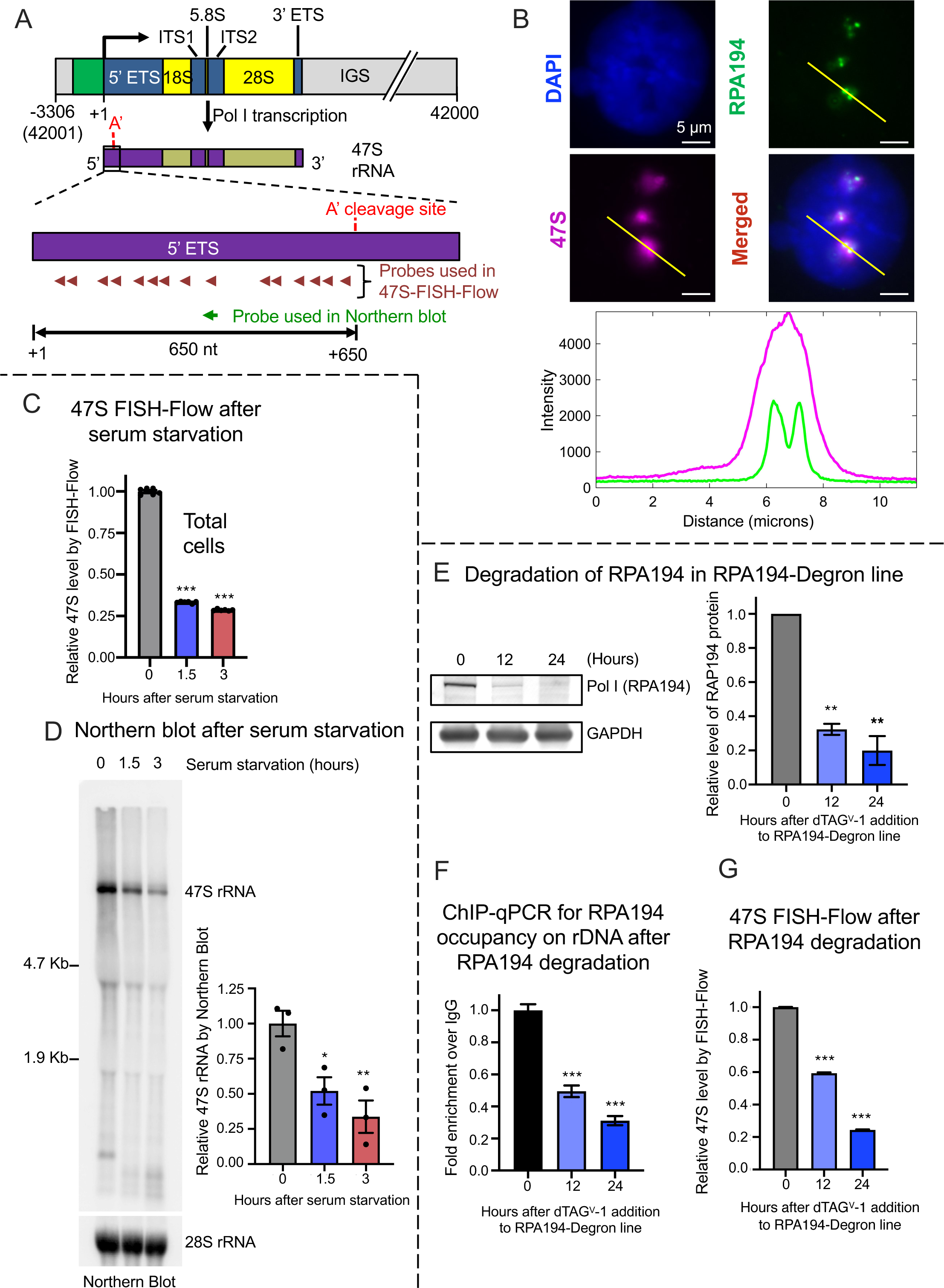
(A) Detailed schematic of 47S-FISH-Flow design: Mouse rDNA is transcribed into 47S rRNA, which undergoes cleavage at the A’ site (in addition to a 3’ cleavage site) to form 45S rRNA. 47S-FISH-Flow utilizes a pool of 15 FISH probes distributed throughout the 650 nucleotide 5’ segment upstream of the A’ cleavage site. Also depicted is the location of the standard Northern blot probe used to quantify 47S. (B) <u>Top</u>) Additional widefield microscopy images showing nucleolar hybridization of 47S-FISH-Flow probes, along with immunofluorescence with anti-RPA194 antibody to mark actively transcribed rDNA alleles, and DAPI for overall DNA. Scale bar (white line) at bottom right indicates 5 µm. B<u>ottom)</u> Fluorescence intensity profile (quantified using ImageJ software) of 47S-FISH-Flow probes (magenta curve) and RPA194 (green curve) plotted across the length of the yellow line drawn in the top images, confirming that 47S FISH-Flow signal, as expected, surrounds Pol I foci. (C) Bargraph showing effect of serum starvation on median 47S rRNA level quantified using 47S-FISH-Flow on the total cell population. n = 6 replicates. (D) <u>Left</u>) Representative northern blot showing level of 47S rRNA in bulk RNA after serum starvation, with mature 28S rRNA shown below as loading control. R<u>ight)</u> Bar graph showing densitometry-based quantification of 47S rRNA normalized to 28S rRNA. n = 3 replicates. It is noted that replicate-to-replicate variability in 47S-FISH-Flow (Fig S6C) is substantially lower than that observed in Northern blot quantification (Fig S6D). (E) <u>Left)</u> Representative western blot showing relative level of Pol I (probed using anti-RPA194) in ‘RPA194-Degron line’ after treatment with 1 μM dTAG^V^-1 for indicated times. The RPA19-Degron line was generated by integrating the FKBPV degron into the endogenous *Polr1a* locus with an identical strategy as used for the *Cebpa* locus in Fig 3A (see Methods). R<u>ight)</u> Bargraph showing relative protein quantification of RPA194 level at 12 and 24 hours. n = 2 replicates. F) Bargraph showing effect of RPA194 degradation on rDNA occupancy: ChIP-qPCR for RPA194 in the RPA194-Degron line treated with 1 μM dTAG^V^-1 for indicated times. It is noted that though dTAG^V^-1 eliminates a majority of total cellular RPA194 by 12 hours (Fig S6E), rDNA-occupied RPA194 is only reduced by 50%, and is resistant to complete degradation. It is plausible that all residual Pol I complexes in the cell at 12 and 24 hours are fully occupied on rDNA. G) Bargraph showing effect of RPA194 degradation on median 47S rRNA level quantified using 47S-FISH-Flow. n = 3 replicates. Magnitude of 47S reduction matches timecourse reduction in RPA194 rDNA occupancy (Fig S6F). All bargraphs show mean plus or minus SEM. *p < .05; **p < .01; ***p < .001, by 2-way Anova with Sidak’s multiple comparison testing. ^56^

**Figure S7 (related to Figure 4).**
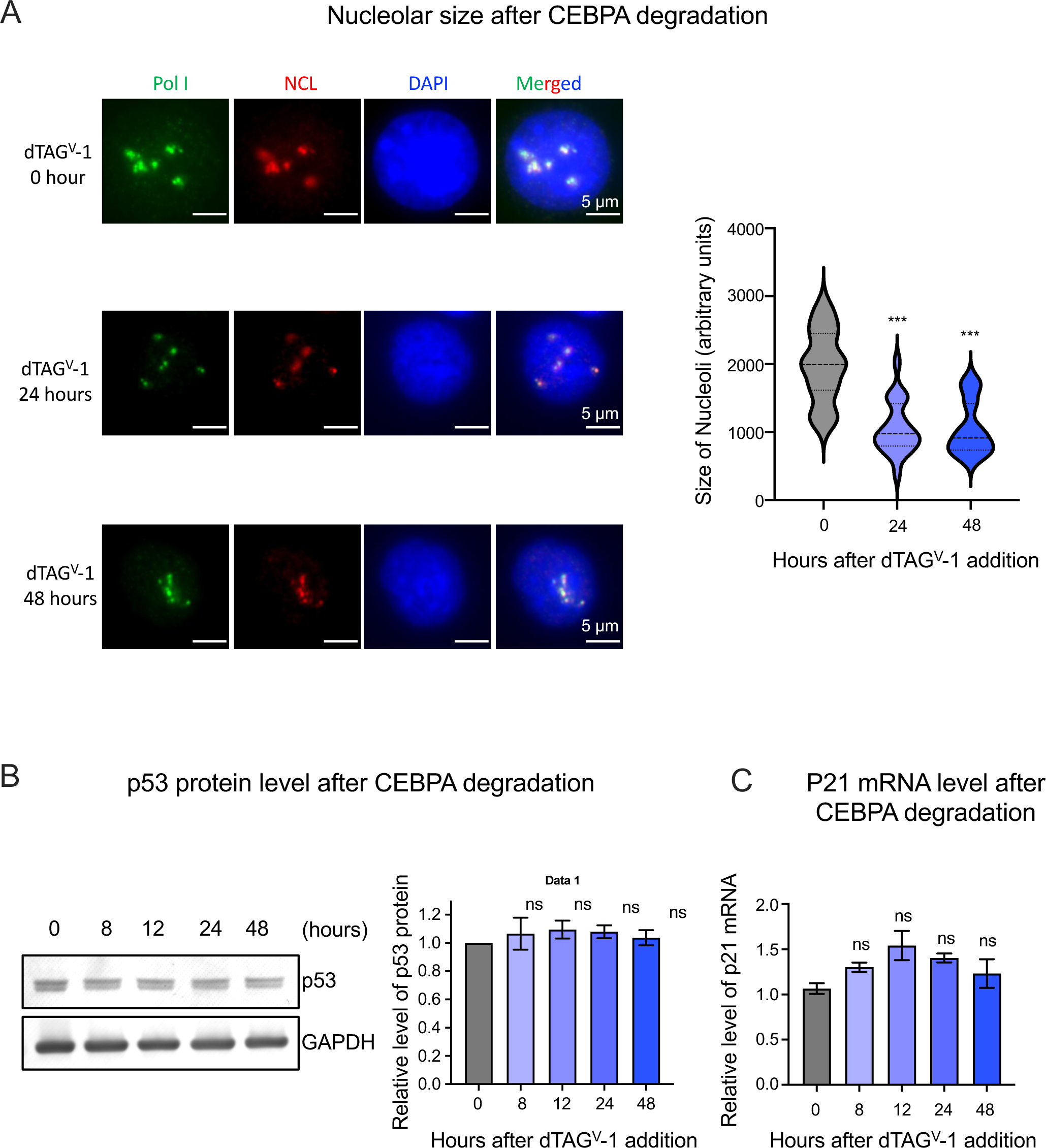
(A) <u>Left</u>) Widefield microscopy images showing nucleoli in the CEBPA-Degron line after treatment with 500 nM dTAG^V^-1 for indicated times. Nucleolar boundary is marked by immunofluorescence with anti-Nucleolin (Red), and actively transcribed rDNA foci with anti-RPA194 (Green), with DAPI to stain overall DNA (Blue). Scale bar (white line) at bottom right indicates 5 µm. R<u>ight)</u> Bargraph showing nucleolar size as quantified by 2D-area-per-cell of Nucleolin signal. n = 20 replicates. (B) <u>Left</u>) Representative western blot showing relative level of p53 protein in CEBPA-Degron line after treatment with 500 nM dTAG^V^-1 for indicated times. R<u>ight)</u> Bargraph showing relative p53 protein quantitation. n = 2 replicates. (C) Bargraph showing RT-qPCR results for p21 mRNA level in CEBPA-Degron line after treatment with 500 nM dTAG^V^-1 for indicated times. n = 2 replicates. All bargraphs show mean plus or minus SEM. *p < .05; **p < .01; ***p < .001, by 2- way Anova with Sidak’s multiple comparison testing.

