## Supplementary material for "Control of Ribosomal RNA Synthesis by Hematopoietic Transcription Factors": Key Resource Table

**Key resources table**

| **REAGENT or RESOURCE** | **SOURCE** | **IDENTIFIER** |
| --- | --- | --- |
| **Antibodies** | | |
| Mouse monoclonal anti-FLAG | Sigma-Aldrich | Cat# F1804; RRID: AB_262044 |
| Rabbit polyclonal anti-CEBP Alpha | Abcam | Cat# ab15048; RRID: AB_2077890 |
| Rabbit monoclonal anti-GAPDH | Cell Signaling Technology | Cat# 2118; RRID: AB_561053 |
| Mouse monoclonal anti-RPA194 (C-1) | Santa Cruz | Cat# sc-48385; RRID: AB_675814 |
| Rabbit polyclonal anti-RRN3 | Proteintech | Cat# 25918-1-AP; RRID: AB_2880296 |
| Rabbit polyclonal anti-SL-1 | (Herdman et al., 2017) | N/A |
| Mouse monoclonal anti-UBTF | Santa Cruz | Cat# sc-13125; RRID: AB_671403 |
| Mouse anti-p53 (DO-1) | Santa Cruz | Cat# sc-126; RRID: AB_628082 |
| Mouse Isotype control | Thermo Fisher Scientific | Cat# 31903; RRID: AB_10959891 |
| Rabbit Isotype control | Thermo Fisher Scientific | Cat# 02-6102; RRID: AB_2532938 |
| Rabbit polyclonal anti-fibrillarin | Abcam | Cat# ab5821; RRID: AB_2105785 |
| Mouse monoclonal anti-Nucleolin, Alexa flour 647 conjugated | Abcam | Cat# ab198580; RRID: AB_2920844 |
| Goat anti-mouse IgG (H+L) antibody, Alexa Fluor 488 conjugated | Molecular Probes | Cat# A-11017; RRID: AB_143160 |
| Donkey anti-rabbit IgG polyclonal antibody, IRDye 680RD conjugated | LI-COR Biosciences | Cat# 925-68073; RRID: AB_2716687 |
| Donkey anti-Mouse IgG (H + L) polyclonal antibody, IRDye 800CW conjugated | LI-COR Biosciences | Cat# 925-32212; RRID: AB_2716622 |
| **Bacterial and virus strains** | | |
| Biological samples |  |  |
| **Chemicals, peptides, and recombinant proteins** | | |
| 3X FLAG peptide | Sigma-Aldrich | Cat#: F4799 |
| β-Estradiol | Fisher Scientific | Cat#: MP021016562 |
| Hanks' Balanced Salt Solution | Thermo Fisher Scientific | Cat#: 14025076 |
| DPBS | GIBCO | Cat#: 14190136 |
| DAPI | Invitrogen | Cat#: D1306 |
| Slow fade Diamond Antifade mounting media | Thermo Scientific | Cat#: S36963 |
| Formaldehyde | Fisher Scientific | Cat#: BP531-500 |
| Formamide | Thermo Scientific | Cat#: AM9342 |
| 20X SSC | Thermo Scientific | Cat#: AM9763 |
| Dextran sulfate | Sigma-Aldrich | Cat#: D8906-50G |
| Cas9 | Thermo Scientific | Cat#: A36498 |
| dTAGV-1 | Tocris | Cat#: 6914 |
| Protease inhibitor cocktail | Sigma-Aldrich | Cat#: P8340 |
| Chymostatin | Cayman Chemical Company | Cat#: 15114 |
| Glycine | Thermo Fisher Scientific | Cat#: 15527013 |
| Micrococcal nuclease | NEB | Cat#: M0247S |
| Sarkosyl | Sigma-Aldrich | Cat#: 61747 |
| Drosophila chromatin | Active Motif | Cat#: 53083 |
| RNase A | Fisher scientific | Cat#: FEREN0531 |
| Dynabeads protein G | Thermo Fisher | Cat#: 10004D |
| anti-H2Av | Active Motif | Cat#: 61686 |
| Proteinase K | Invitrogen | Cat#: 25530049 |
| 2X Luna universal qPCR master mix | NEB | Cat#: M3003X |
| DNase I | Thermo Fisher | Cat#:18068015 |
| Ribonucleoside vanadyl complex | NEB | Cat#: S1402S |
| Poly-D-lysine | Nalgene | Cat#: 343910001 |
| Rectangular cover glass, #1.5 thickness | Thomas Scientific | Cat#: 1217N86 |
| Serum | Cell Signaling Technology | Cat#: 5425 |
| Hybond-N membrane | Cytiva Life Sciences | Cat#: RPN303N |
| ULTRAhyb oligo buffer | Thermo Fisher Scientific | Cat#: AM8663 |
| [γ-32P] ATP | PerkinElmer | Cat#: BLU502A001MC |
| T4 polynucleotide kinase | NEB | Cat#: M0201S |
| **Critical commercial assays** | | |
| Neon transfection kit | Thermo Fisher Scientific | Cat#: MPK1025 |
| BCA protein assay kit | Thermo Fisher Scientific | Cat#: 23227 |
| Qiagen PCR purification kit | Qiagen | Cat#: 28106 |
| NEBNext Ultra II DNA library prep kit | NEB | Cat#: E7645S |
| NEBNext Multiplex Oligos for Illumina | NEB | Cat#: E7600S |
| KAPA library quantification kit | Fisher scientific | Cat#: KK4824 |
| Qiagen RNeasy Mini Kit | Qiagen | Cat#: 74104 |
| Primescript RT reagent kit | Takara Bio | Cat#: RR037A |
| **Deposited data** | | |
| Raw western blot and microscopy data | This paper | Mendeley: <https://data.mendeley.com/datasets/kfp6hf7c2s/draft?a=90d1efff-6a42-49db-aebb-d8578b9abb05> |
| TF-rDNA Atlas analyzed data | This paper | GEO: GSE193651 |
| ChIP-Seq studies in CEBPA-Degron line | This paper | GEO: GSE193651 |
| **Experimental models: Cell lines** | | |
| ER-HoxA9 myeloid GMP cell line | (Sykes et al., 2016) |  |
| Chinese hamster ovary cell line |  |  |
| **Experimental models: Organisms/strains** | | |
| **Oligonucleotides** | | |
| SgRNAs, PCR Primers, FISH-Flow probes, and Northern blot probe are listed in Table S3 | This paper | N/A |
| **Recombinant DNA** | | |
| pUC57-CEBPA homology-FKBPV-FLAG | This paper | N/A |
| pUC57-RPA194 homology-FKBPV-FLAG | This paper | N/A |
| **Software and algorithms** | | |
| Stellaris-probe-designer | Biosearch technologies | <https://www.biosearchtech.com/support/tools/design-software/stellaris-probe-designer> |
| SgRNAs Design Tool | CHOPCHOP | <https://chopchop.cbu.uib.no/> |
| GraphPad Prism (version 9.4.0) | GraphPad Software Inc | <https://www.graphpad.com/scientific-software/prism/> |
| FlowJo (version 10.8.1) | FlowJo, LLC | <https://www.flowjo.com/solutions/flowjo> |
| Fiji/ImageJ | Fiji/ImageJ | <https://imagej.net/software/fiji/> |
| ImageQuant software | Cytiva life sciences | <https://www.cytivalifesciences.com/en/us/shop/molecular-biology/nucleic-acid-electrophoresis--blotting--and-detection/molecular-imaging-for-nucleic-acids/imagequant-tl-10-1-analysis-software-p-28619> |
| Bowtie2 | (Langmead and Salzberg, 2012) | <http://bowtie-bio.sourceforge.net/bowtie2/index.shtml> |
| Samtools | MIT/Expat | <http://www.htslib.org/> |
| IGVtools | Broad institute | <https://software.broadinstitute.org/software/igv/igvtools> |
| R software | The R Foundation | <https://www.r-project.org/> |
| Image Studio version 5.2.5 | LI-COR | <http://opensource.licor.com/licenses/ImageStudio/index.html> |
| **Other** | | |
